# Astrocytic glycogenolysis gates Warburg-like metabolic reprogramming that promotes neuropathic pain chronification

**DOI:** 10.1101/2025.05.27.655245

**Authors:** Jun Seo Park, Kwang Hwan Kim, Hye Won Jun, Se-Young Choi, Seung Bum Park, Sung Joong Lee

## Abstract

Chronic pain remains a major unmet medical challenge, yet the metabolic checkpoints that govern its persistence are poorly defined. Astrocytes are increasingly recognized as chemically programmable hubs that tune neuronal excitability through metabolic circuits. Building on reports that astrocyte–neuron lactate shuttling (ANLS) in the anterior cingulate cortex (ACC) supports chronic pain, we asked how astrocytic metabolic states evolve over the course of pain chronification. Using untargeted metabolomics of the ACC combined with GFAP-RiboTag–based astrocyte-specific transcriptomics, we provide a time-resolved map of astrocytic metabolism across the transition from acute nociception to chronic neuropathic pain. This analysis reveals a biphasic glycogen program—an acute glycogenolysis-triggered glycogen supercompensation—that culminates in the emergence of a Warburg-like metabolic signature with tissue acidification associated with sustained lactate shuttling and persistent circuit activation. Using glycogen phosphorylase inhibitors (GPI-1, GPI-2) as pharmacological probes, we show that early glycogenolysis blockade attenuates this Warburg-like shift, partially normalizes ACC metabolic signatures, and reduces long-lasting mechanical hypersensitivity, without impairing acute nociceptive sensitization. These findings identify astrocytic metabolic reprogramming as a pharmacologically tractable circuit-level process and nominate glycogenolysis as an upstream biochemical gate and potential therapeutic control point in neuropathic pain.

**HIGHLIGHTS:** Most experimental models of chronic pain emphasize progressive sensitization, yet few have identified a chemically tractable switch that converts transient nociception into a persistent cortical state. Here, we propose astrocytic glycogenolysis as such a switch—a gatekeeper process whose brief engagement can reconfigure neuroglial metabolism toward a Warburg-like signature that stabilizes hyperexcitable circuit dynamics. By leveraging small-molecule glycogen phosphorylase inhibitors as mechanistic precision probes, we move beyond correlation to delineate a pathway-level control architecture that can be transiently perturbed to dissociate acute nociceptive processing from pain chronification. This framework reframes astrocytes from metabolic support cells to state-setting regulators of cortical excitability and positions glycogenolysis as a therapeutically actionable leverage point. More broadly, our findings underscore a methodological opportunity: applying chemical biology to neuroglial metabolism can expose governing checkpoints that are often obscured by descriptive physiology, enabling mechanism-first, pathway-directed intervention strategies across neurological disease contexts.

## INTRODUCTION

Neuropathic pain is a persistent pathological condition that arises from injury or dysfunction of the nervous system, in which repeated or intense nociceptive input drives the transition from acute to enduring pain.^[1,2]^ Among cortical regions, the anterior cingulate cortex (ACC) has been implicated as a key hub for pain chronification.^[3,4]^ Persistent hyperexcitability of the ACC is a key contributor to the maintenance of chronic pain networks and identifies ACC excitability as a representative circuit-level target whose modulation can alleviate pain-related phenotypes.^[3,4]^

Astrocytes are key modulators of brain function, shaping neuronal excitability and circuit dynamics through metabolic interactions with neurons.^[5,6]^ They are increasingly recognized as metabolic “switchboards” that integrate neuromodulatory, inflammatory, and energetic cues.^[6–9]^ This metabolic control is governed by metabolite fluxes and enzymatic checkpoints, which can be selectively probed using small-molecule tools.^[10,11]^ In particular, the astrocyte–neuron lactate shuttle (ANLS) has long been recognized as a central mechanism by which astrocytes couple their metabolism to that of neurons by supplying lactate as both an energy substrate and a signaling cue.^[6,7,8,12]^ Astrocyte-derived lactate release to neurons has been reported to modulate neuronal energy metabolism, gene expression, and synaptic plasticity, thereby reconfiguring long-term circuit activity and behavior.^[12–22]^

In neuropathic pain models, expression of ANLS components and lactate shuttling within the anterior cingulate cortex (ACC) are increased, and pharmacological blockade of astrocyte-derived lactate transport reduces pain behaviors, positioning ANLS as a key metabolic axis in pain chronification.^[23–25]^ However, prior ACC studies have largely relied on end-point measurements of bulk lactate levels, astrocytic Ca^2^⁺ signaling, or generalized reactive gliosis.^[18,19,23–25]^ Nevertheless, the temporal trajectory followed by ACC ANLS during the transition from acute nociceptive input to a chronic pain state, and the upstream metabolic signals that drive this reorganization, remain incompletely defined.

Within the ANLS, lactate supplied to neurons is generated primarily via astrocytic glycolysis, and this glycolytic flux is thought to depend critically on glucose derivatives provided by glycogen breakdown within astrocytes.^[21,23,26–28]^ However, despite repeated observations linking glycogen levels to lactate production, it remains largely unresolved how astrocytic glycogen metabolism is temporally reprogrammed over the course of chronic pain, and through which intermediate metabolic programs this reconfiguration acts to regulate ANLS.

Conceptually, enhanced lactate metabolism in the neuropathic pain parallels the Warburg effect described in cancer.^[29,30]^ The Warburg effect is characterized by increased glycolytic flux in the presence of sufficient oxygen, accompanied by lactate accumulation and tissue acidification.^[30,31]^ Importantly, it reflects not only increased lactate production but also a coordinated metabolic reprogramming that propagates into diverse signaling changes.^[29–31]^ Such Warburg-like features are increasingly appreciated as hallmarks of pathological states, including certain forms of immune activation and chronic inflammation.^[29–31]^ Although altered energy metabolism and elevated lactate have been reported in pain-related circuits, whether astrocytes in the ACC undergo a broader Warburg-like reprogramming—and how such a state would relate to glycogen metabolism and ANLS—remain essentially unknown. This raises the broader question of how astrocytic glycogenolysis, lactate shuttling, and potential Warburg-like metabolic states are temporally organized during neuropathic pain.

Here, we asked how astrocytic metabolic states evolve across the time course of pain chronification and what upstream “gating” signals orchestrate this transition. Using GFAP-RiboTag–based astrocyte-specific transcriptomic profiling together with untargeted metabolomics, we mapped the first time-resolved astrocytic metabolic remodeling during pain progression. We find that a Warburg-like metabolic signature emerges as pain becomes chronic and that this program is gated by an acute bout of glycogenolysis that is followed by glycogen supercompensation. These findings connect classical brain glycogen physiology to ACC pain circuits and delineate a mechanistic link between acute nociception and persistent circuit activation underlying pain chronification, suggesting metabolic nodes that could be targeted for intervention.

## RESULTS

### Astrocyte–neuron lactate shuttling (ANLS) displays distinct temporal dynamics

Across diverse chronic pain models (e.g., inflammatory, visceral, and neuropathic), chronic pain states are associated with elevated lactate levels in the ACC, and astrocyte-derived lactate release through the ANLS has been implicated in pain modulation.^[23–25]^ However, most of these studies have focused on single time points, and how lactate- and ANLS-related processes are initiated, coordinated, and maintained over the full course of pain chronification remains poorly defined.^[23–25]^

To address this knowledge gap, we sought to characterize the temporal profile of ANLS in the ACC during the transition from acute to chronic pain. Neuropathic chronic pain was induced using the spinal nerve transection (SNT) model (Figure 1A).^[32–34]^ Compared with Sham controls, SNT mice exhibited prolonged mechanical allodynia, confirming the development of chronic pain behavior (Figure 1B). We next generated GFAP-RiboTag mice to capture astrocyte-specific changes in gene expression (Figure 1A and Figure S1). The generation of GFAP-RiboTag mice was validated by PCR and Western blotting (Figure S1A,B). In GFAP-RiboTag mice subjected to SNT, ACC tissue was collected at three time points (day 1, day 3, and day 7 after surgery), and transcriptomic analyses were performed to delineate the temporal dynamics of astrocytic ANLS-related gene expression. qPCR analyses revealed that genes regulating lactate production (lactate dehydrogenase A/B, Ldha/b) and transport (monocarboxylate transporters, Mcts) exhibited biphasic dynamics: both were suppressed at day 3 but robustly upregulated in the chronic state (day 7) (Figures 1C–F). These results, consistent with prior work implicating ANLS in chronic pain,^[23–25]^ suggest that ANLS activity is not continuously elevated but rather gated in a time-dependent manner.

**Figure 1.**
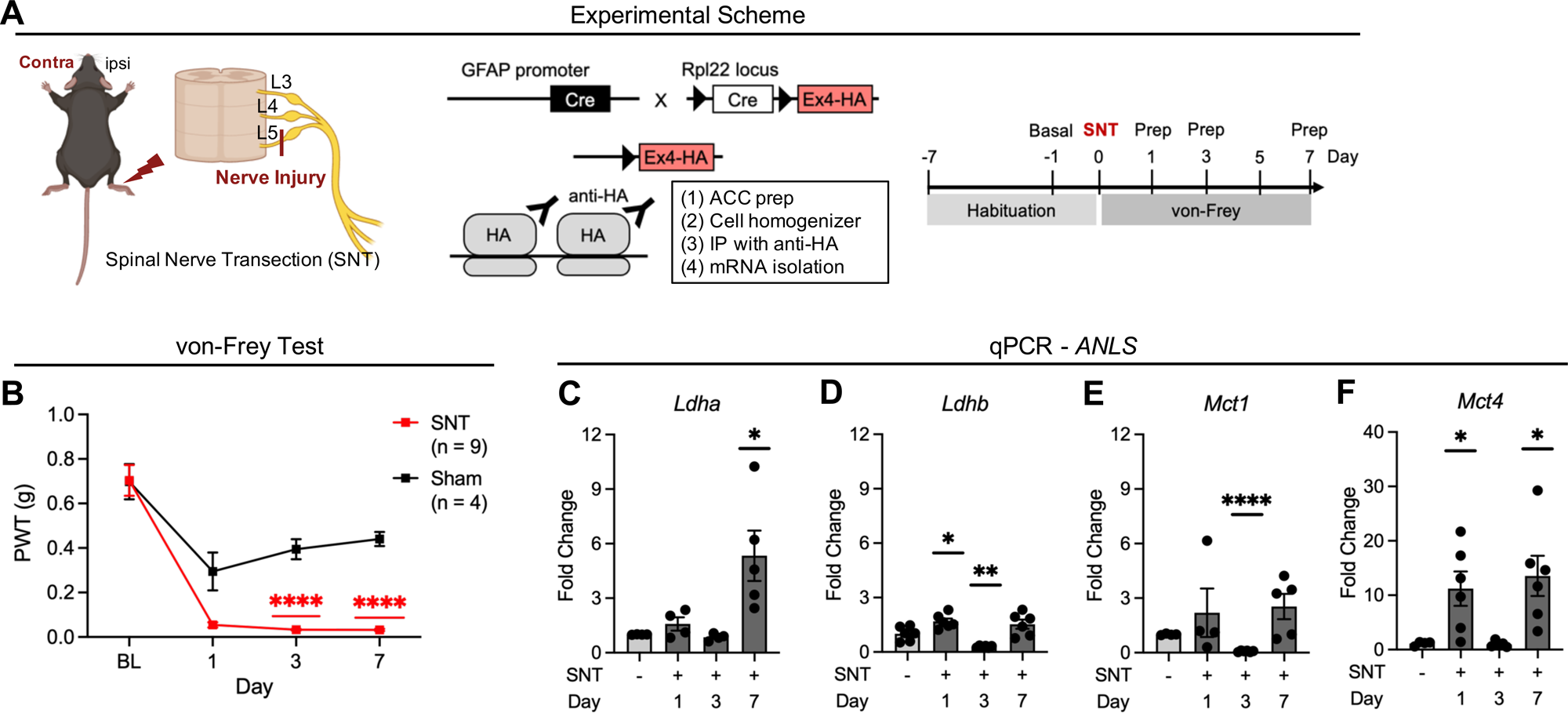
Astrocyte–neuron lactate shuttling displays distinct temporal dynamics in neuropathic pain. (A) Schematic illustration of neuropathic pain surgery and GFAPCre::RiboTag mice transcriptomics experiment. (B) von-Frey test of neuropathic pain model. ****P < 0.0001 (3 day SNT vs 3 day Sham), ****P < 0.0001 (7 day SNT vs 7 day Sham) (C-F) qPCR data of ANLS gene profiles. (Time points: 1 day, 3 day and 7 day), (n = 4~6) (C) Fold change of *Ldha* in each time point. *P = 0.0351 (7 day SNT vs Sham) (D) Fold change of *Ldhb* in each time point. *P = 0.0136 (1 day SNT vs Sham), **P = 0.0064 (3 day SNT vs Sham) (E) Fold change of *Mct1* in each time point. ****P < 0.0001 (3 day SNT vs Sham) (F) Fold change of *Mct4* in each time point. *P = 0.0243 (1 day SNT vs Sham), *P = 0.0199 (7 day SNT vs Sham) Data are represented as the mean ± SEM; *P < 0.05, **P < 0.01, ***P < 0.001, ****P < 0.0001; Two-way ANOVA followed by Bonferroni’s multiple comparisons test (B) and Student’s t-test (C-F).

### Astrocyte Warburg-like metabolic reprogramming in pain chronification

We observed that astrocytic responses during pain chronification follow distinct temporal trajectories. To extend these observations, we conducted time-resolved metabolomic analyses to delineate how metabolic profiles are dynamically reconfigured during the transition from acute to chronic pain.^[35]^ We performed untargeted bulk metabolomics on the contralateral ACC samples collected at 0.5, 1, 3, and 7 days after SNT using GC-MS (Figures 2A and S2). All metabolite data were first normalized to the sample values from the ipsilateral region and subsequently expressed as fold changes relative to Sham controls (n = 4) and subsequently subjected to statistical analysis to evaluate significance (Figure S2).^[36,37]^ Both principal component analysis (PCA) and partial least squares-discriminant analysis (PLS-DA) revealed progressive separation of metabolic profiles from the acute state (day 0.5, 1) to the chronic state (day 7), with strong loadings on lactate, α-ketoglutarate, and 3-phosphoglycerate (3-PG) (Figure 2B,C). At the single-metabolite level, several species displayed time-dependent increases. Specifically, glycolytic intermediates (lactate and 3-phosphoglycerate) and several amino acids (L-glutamate and L-glutamine) increased across time points, whereas GABA (4-aminobutanoic acid) decreased; most fatty acids showed no significant change (Figure 2D).

**Figure 2.**
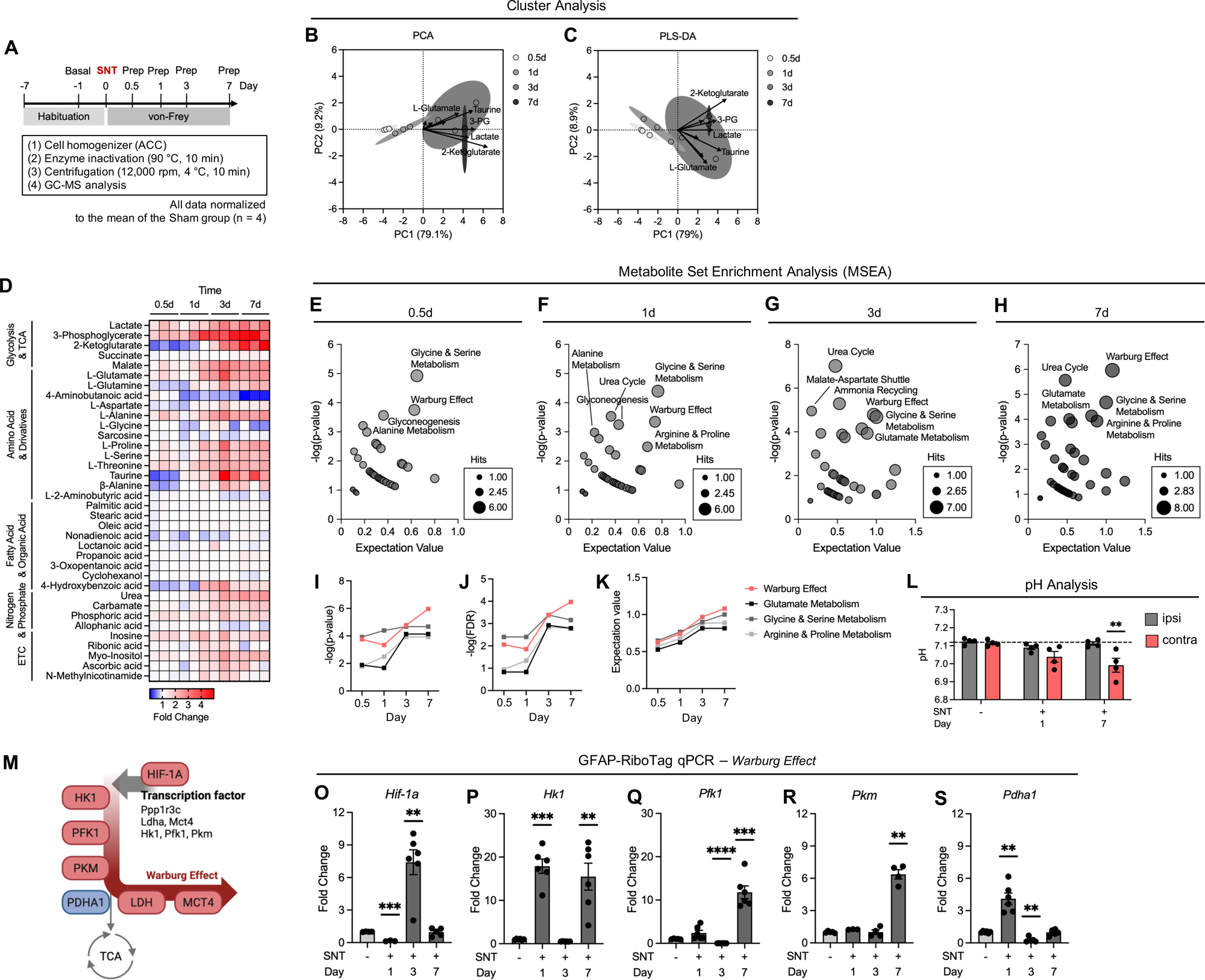
Astrocyte Warburg-like metabolic reprogramming in pain chronification. (A) Schematic illustration of GC-MS metabolomics analysis in SNT pain model. (B and C) Cluster analysis of GC-MS data; compared by each time point (0.5d, 1d, 3d, 7d). Show axis of representative metabolite. Arrows indicate loading vectors for representative metabolites (lactate, 3-phosphoglycerate(3-PG), 2-ketoglutarate L-glutamate and taurine). (Additional statistical analyses and supporting evidence are provided in the Supplementary Information; validated by cross-validation and permutation testing.) (B) Principal component analysis (PCA) of GC-MS fold-change profiles of each time point. Cluster separation was observed in acute time points (0.5d, 1d) versus chronic time point (7d). The variance in X-axis (PC1) is 79.1% and on Y-axis (PC2) is 9.2%. (C) Partial least squares-discriminant analysis (PLS-DA) of GC-MS fold-change profiles of each time point. Cluster separation was observed in acute time points (0.5d, 1d) versus chronic time point (7d). The variance in X-axis (PC1) is 79% and on Y-axis (PC2) is 8.9%. (D) Heatmap of 28 metabolites constant fold-change in each time point (Time points: 0.5 day(12h), 1 day, 3 day and 7 day), (n = 3). (E-H) Metabolite set enrichment analysis (MSEA) of GC-MS data; compared by each time point (0.5 day, 1 day, 3 day and 7 day). Labeling selected high enrichment metabolic pathway. (E) 0.5 day bubble plot of after SNT. (F) 1 day bubble plot of after SNT. (G) 3 day bubble plot of after SNT. (H) 7 day bubble plot of after SNT. (I-K) Analysis of the time-dependent enrichment of selected pathways. (Time points: 0.5 day, 1 day, 3 day and 7 day) (I) −log(p-value) of time-dependent enrichment of selected pathways. (J) −log(FDR) of time-dependent enrichment of selected pathways. (K) Expectation value of time-dependent enrichment of selected pathways. (L) pH of ACC in each time point (Time points: 1 day and 7 day), \*\**P* = 0.0096 (SNT 7 day contra vs Sham), (n = 4); Two-way ANOVA-multiple comparisons (M) Schematic illustration of Warburg effect pathway. (Created by https://www.biorender.com) (O-S) qPCR data of Warburg effect signiture gene profiles. (Time points: 1 day, 3 day and 7 day), (n = 4~6) (O) Fold change of *Hif-1a* in each time point. ***P = 0.0002 (1 day SNT vs Sham), **P = 0.0025 (3 day SNT vs Sham) (P) Fold change of *Hk1* in each time point. ***P = 0.0002 (1 day SNT vs Sham), **P = 0.0055 (7 day SNT vs Sham) (Q) Fold change of *Pfk1* in each time point. ****P < 0.0001 (3 day SNT vs Sham), ***P = 0.0002 (7 day SNT vs Sham) (R) Fold change of *Pkm* in each time point. **P = 0.0011 (7 day SNT vs Sham) (S) Fold change of *Pdha1* in each time point. **P = 0.0019 (1 day SNT vs Sham), **P = 0.0013(3 day SNT vs Sham) Data are represented as the mean ± SEM; *P < 0.05, **P < 0.01, ***P < 0.001, ****P < 0.0001; Student’s t-test (O-S).

Accordingly, we performed pathway enrichment analyses to interrogate the organization of metabolic pathways and observed that the enrichment profiles varied across the different time points.^[36,37]^ Distinct metabolic pathways were progressively reinforced across stages, with the Warburg effect signiture exhibiting the most prominent enrichment in the chronic (day 7) state (Figures 2E-H). Pathway enrichment increased over time for the Warburg effect signiture and selected pathways (e.g., glutamate metabolism), with progressively stronger statistical support, as indicated by rising −log10(p), −log10(FDR), and expectation values (Figure 2I-K). Consistently, gene set enrichment analysis (GSEA) of ACC bulk-transcriptome data at day 7 in the chronic pain model revealed upregulation of Warburg–effect–associated and glutamate metabolism signatures. In contrast, the tricarboxylic acid (TCA) cycle and oxidative phosphorylation (OXPHOS) programs were downregulated (Figure S3). The Warburg effect—traditionally associated with cancer and hypoxic conditions—characterizes glycolytic reprogramming, accompanied by elevated lactate production.^[29,30]^ Consistent with this signature, tissue pH was significantly reduced in the contralateral ACC during the chronic state compared with Sham or the ipsilateral controls (Figure 2L).

To determine whether astrocytes themselves engage this program, we used GFAP-RiboTag profiling of canonical Warburg genes. The transcription factor hypoxia-inducible factor-1α (HIF-1α)—a master regulator of the Warburg phenotype by promoting lactate production (via *Ldha*) while limiting pyruvate flux into the TCA cycle (Figure 2N)—was robustly expressed at the intermediate (day 3) time point (Figure 2O). Its downstream targets, including hexokinase 1 (*Hk1*), phosphofructokinase 1 (*Pfk1*), and pyruvate kinase M1/M2 (*Pkm*) which inducing a metabolic shift toward lactate production, were consistently strongly upregulated by 7 days (Figures 2P-R), paralleling increases in *Ldha* and *Mct4* (Figures 1C and 1F). In contrast, pyruvate dehydrogenase E1α (*Pdha1*), which directs pyruvate into the TCA cycle, was reduced (Figure 2S).

Together, these data extend the ANLS model of chronic pain to a Warburg-like metabolic reprogramming. We observe transcriptional priming at intermediate stages, followed by full engagement of a Warburg program at 7 days, culminating in elevated astrocytic lactate production.

### Astrocyte glycogen supercompensation in pain chronification

Not only in pain paradigms but also across diverse experimental models, ANLS has been implicated as an important mechanism in regulating neuronal activity and behavior.^[6,12]^ Within this framework, glycogen breakdown (glycogenolysis) has repeatedly been proposed as an upstream metabolic pathway that controls ANLS.^[13,14,21,26–28]^ Given this, we sought to determine how the glycogen metabolic program is remodeled over pain chronification and, therefore, focused on glycogen metabolism as an upstream control node.

Glycogen flux is bidirectionally regulated: synthesis is mediated by *Gys1* activation downstream of *Ppp1r3c*, while breakdown is mediated by *Pygb* activation downstream of *Phka2* (Figure 3A). To test whether these dynamics are altered in vivo, we assessed astrocyte gene expression in RiboTag mice (Figure 3B).

**Figure 3.**
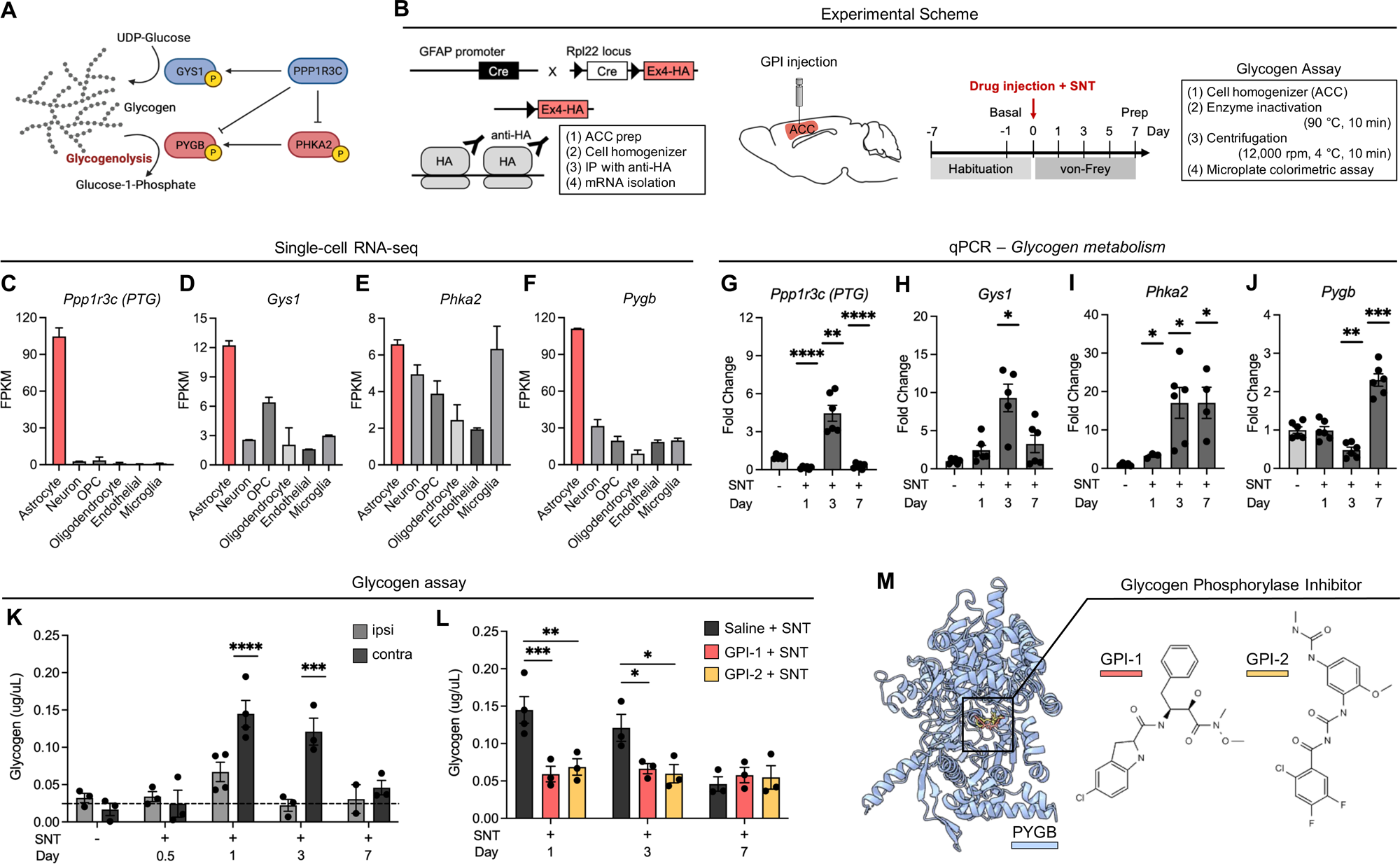
Astrocyte glycogen supercompensation in pain chronification. (A) Scheme of glycogen metabolism pathway. (Created in https://www.biorender.com) (B) Schematic illustration of GFAPCre::RiboTag mice and neuropathic pain model transcriptomics experiment (left). Schematic illustration of Glycogen assay and GPI-1 stereotaxic injection in ACC (rignt). (C-F) Glycogen metabolism gene (*Ppp1r3c, Gys1, Phka2, Pygb*) single cell RNA-sequencing expression result of each cell type. (C) Single cell RNA seq data of *Ppp1r3c.* (D) Single cell RNA seq data of *Gys1.* (E) Single cell RNA seq data of *Phka2.* (F) Single cell RNA seq data of *Pygb.* (G-J) qPCR data of glycogen metabolism gene (*Ppp1r3c, Gys1, Phka2, Pygb*). (Time points: 1d, 3d and 7d), (n = 4~6), (GFAP-RiboTag). (G) Fold change of *Ppp1r3c* in each time point. ****P < 0.0001 (1 day SNT vs Sham), **P = 0.0026 (3 day SNT vs Sham), ****P < 0.0001 (7 day SNT vs Sham). (H) Fold change of *Gys1* in each time point. *P = 0.0101 (3 day SNT vs Sham). (I) Fold change of *Phka2* in each time point. *P = 0.0133 (1 day SNT vs Sham), *P = 0.0104 (3 day SNT vs Sham), *P = 0.0288 (7 day SNT vs Sham). (J) Fold change of *Pygb* in each time point. **P = 0.0011 (3 day SNT vs Sham), ***P = 0.0001 (7 day SNT vs Sham). (K-L) Glycogen assay analysis of ACC, (n=3~4). (K) Glycogen constant in ACC, compare by contralateral area and ipsilateral area. (Time point: Sham, 0.5d, 1d, 3d, 7d) ****P < 0.0001 (1d SNT contra vs ipsi), ***P = 0.0006 (3d SNT contra vs ipsi). (L) Glycogen contant in ACC, compare by contralateral ACC GPI injection. ***P = 0.0005 (1d Saline + SNT vs GPI-1 + SNT), **P = 0.0015 (1d Saline + SNT vs GPI-2 + SNT), *P = 0.0308 (3d Saline + SNT vs GPI-1 + SNT), *P = 0.0148 (3d Saline + SNT vs GPI-2 + SNT). (M) PYGB-Glycogen Phosphorylase Inhibitor(GPI) complex structure and GPI chemical structure of GPI-1 (left) and GPI-2 (rignt). Data are represented as the mean ± SEM; \**P* < 0.05, **P < 0.01, \*\*\**P* < 0.001, \*\*\*\**P* < 0.0001; Student’s t-value test (G-J) and Two-way ANOVA followed by Bonferroni’s multiple comparisons test (K-L).

In the CNS, glycogen metabolism is generally considered to be predominantly enriched in astrocytes.^[26,28]^ As expected, glycogen metabolism was confirmed to be astrocyte-enriched based on cortical single-cell RNA-seq dataset,^[39]^ with enzymes governing both synthesis and breakdown predominantly localized to astrocytes (Figures 3C–F).

Transcriptomic profiling revealed a biphasic program: from the acute (day 1) to the intermediate (day 3) stages, synthesis-side genes *Ppp1r3c* and *Gys1* were elevated, peaking at day 3 (Figures 3G–H). From the intermediate (day 3) to the chronic (day 7) stages, degradation-side genes *Phka2* and *Pygb* were strongly induced (Figures 3I–J). Thus, glycogen metabolism shifts from a synthesis-dominant to a degradation-dominant program, potentially consistent with transcriptional priming by *HIF-1α*, which is known to induce *Ppp1r3c* and aligns with the observed temporal pattern (Figure 3G). Direct measurement of glycogen content in the ACC confirmed this biphasic trajectory. The contralateral ACC exhibited early glycogen accumulation followed by pronounced breakdown; notably, glycogen levels at the day 3 (intermediate stage) were significantly higher than in Sham or ipsilateral controls (Figure 3K). Direct measurement of glycogen content in the ACC confirmed this biphasic trajectory. The contralateral ACC exhibited rapid early glycogen accumulation followed by pronounced breakdown; notably, glycogen levels at the intermediate stage were significantly higher than in Sham or ipsilateral controls (Figure 3K).

The observed pattern of excessive glycogen synthesis followed by breakdown is highly consistent with the classical phenomenon of glycogen supercompensation.^[40,41]^ In cell types with active glycogen metabolism, including astrocytes, an initial phase of glycogenolysis can serve as a trigger for glycogen supercompensation, such that inhibition of this early glycogenolytic phase attenuates the subsequent remodeling of glycogen metabolism.^[28,40,41]^ We therefore interpret the pronounced glycogen accumulation at day 1–3 as a supercompensation peak likely triggered by an earlier glycogenolytic bout initiated immediately after SNT. To confirm whether the initial glycogenolysis led to the glycogen supercompensation during pain chronification, we inhibited acute glycogenolysis at the onset of SNT pain. We first selected a PYGB inhibitor, a small-molecule glycogen phosphorylase inhibitor (GPI), to suppress glycogenolysis.^[42,43]^ Among six glycogen phosphorylase inhibitors previously reported to target the PYGB binding pocket, we prioritized two compounds with the most favorable docking predictions (Figure S4) ^[44–46]^ and subsequently validated their PYGB selectivity by profiling binding across glycogen phosphorylase isoforms and performing reverse target screening (Figure S5).^[47–49]^

Then, we applied each of the two selected GPIs to the ACC and monitored the resulting changes in glycogen dynamics (Figure 3B, 3L–M). In both GPI groups, excessive glycogen accumulation on the first day of SNT was rescued (Figure 3L), supporting the view that early glycogenolysis is required for the subsequent supercompensation-like trajectory during pain chronification. Importantly, this phenotype is abolished by acute suppression of glycogenolysis, identifying glycogen metabolism as a chemically tractable gate for metabolic reprogramming during pain chronification.

### Glycogenolysis inhibition quenches Warburg-like metabolic reprogramming

A mechanistic link between glycogen metabolism and the Warburg effect has been well established, where glycogen supercompensation amplify glycolytic reprogramming.^[52–56]^ To test whether this mechanism underlies pain chronification, we injected GPI-1 or GPI-2 into contralateral ACC immediately after SNT, and performed untargeted NMR metabolomics at the chronic (day 7) stage (Figures 4A and S7).

**Figure 4.**
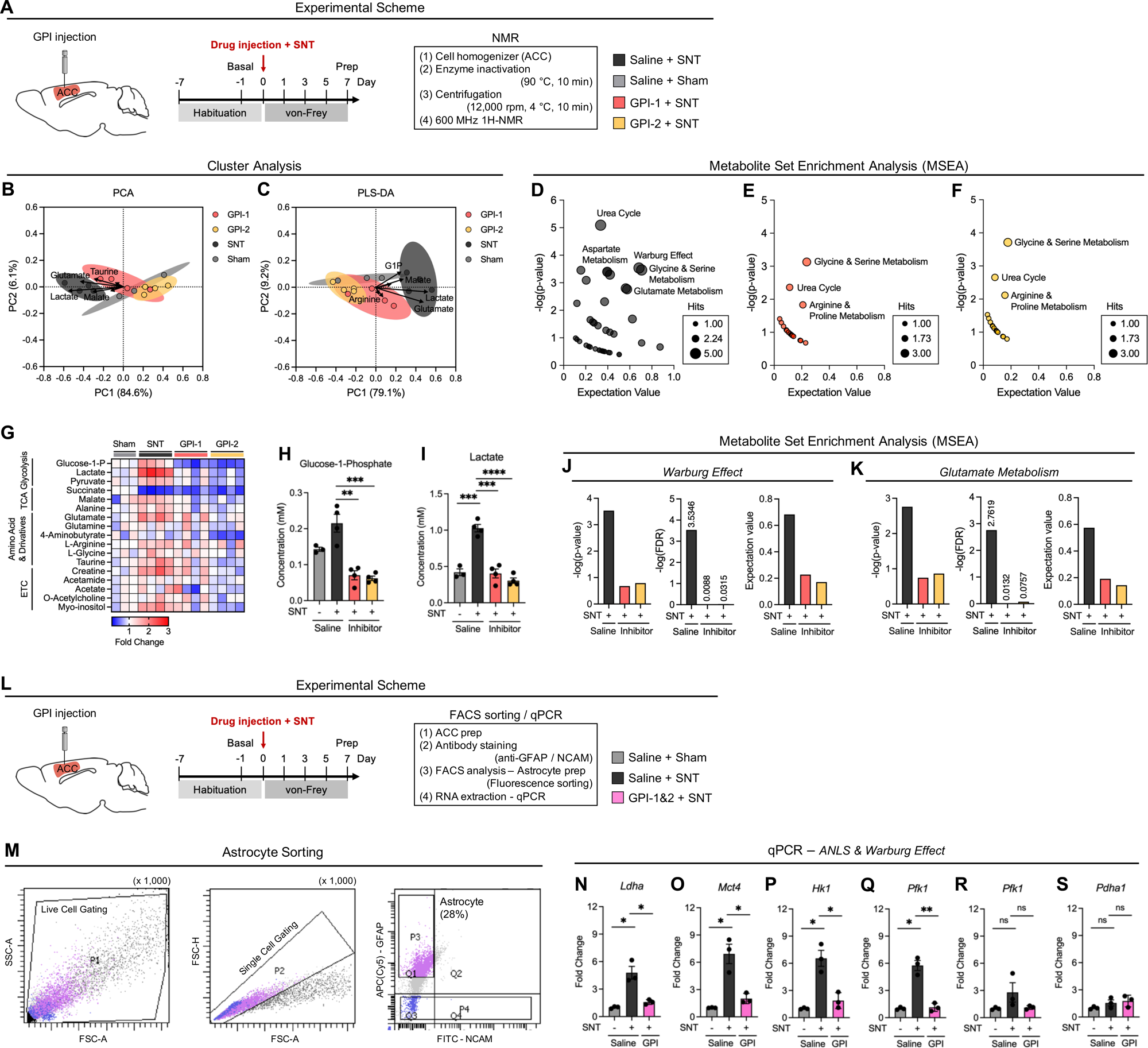
Glycogenolysis inhibition quenches Warburg-like metabolic reprogramming. (A) Schematic illustration of NMR after GPI stereotaxic injection in ACC. (B and C) Cluster analysis of NMR data; compared by each group (GPI-1, GPI-2, SNT, Sham), (n = 4). Arrows indicate loading vectors for representative metabolites (Glucose-1-phosphate (G1P), Lactate, Arginine, Glutamate, Malate and Taurine). (Additional statistical analyses and supporting evidence are provided in the Supplementary Information; validated by cross-validation and permutation testing.) (B) Principal component analysis (PCA) of NMR fold-change profiles of each time point. Cluster separation was observed in SNT group (Saline + SNT) vs GPI injection groups (GPI-1 + SNT, GPI-2 + SNT). The variance in X-axis (PC1) is 84.6% and on Y-axis (PC2) is 6.1%. (C) Partial least squares-discriminant analysis (PLS-DA) of NMR fold-change profiles of each time point. Cluster separation was observed in SNT group (Saline + SNT) vs GPI injection groups (GPI-1 + SNT, GPI-2 + SNT). The variance in X-axis (PC1) is 79.1% and on Y-axis (PC2) is 9.2%. (D-F) Metabolite set enrichment analysis (MSEA) of NMR metabolomics data. Labeling selected high enrichment metabolic pathway. (D) Bubble plot of SNT after saline injection. (E) Bubble plot of SNT after GPI-1 injection. (F) Bubble plot of SNT after GPI-2 injection. (G) Heatmap of 17 metabolites constant fold-change in each group (Group: Saline + SNT, Saline + Sham, GPI-1 + SNT, GPI-2 + SNT), (n = 3~4). (H and I) Metabolite concentration data by NMR analysis. (H) G1P concentration of each group. **P = 0.0020 (GPI-1 + SNT vs Saline + SNT), ***P = 0.0010 GPI-2 + SNT verse Saline + SNT) (I) Lactate concentration of each group. ***P = 0.0003 (Saline + Sham vs Saline + SNT), ***P = 0.0003 (GPI-1 + SNT vs Saline + SNT), ****P < 0.0001 (GPI-2 + SNT vs Saline + SNT) (J and K), Metabolite set enrichment analysis (MSEA) of NMR metabolomics data. Labeling selected high enrichment metabolic pathway (Statistic factor: −log(p-value), −log(FDR), Expectation value). (J) Warburg effect pathway. (K) Glutamate metabolism pathway. (L) Schematic illustration of astrocyte FACS sorting after GPI-cocktail stereotaxic injection in ACC. (M) Representative images of astrocyte sorting strategy. Live cell gate (P1) on FSC-A vs SSC-A to exclude debris (Left). Singlet gate (P2) on FSC-H vs FSC-A (Middle). Bivariate plot of APC(Cy5)-GFAP versus FITC-NCAM used to define astrocytes (GFAP⁺/NCAM⁻; gate P3; representative frequency ~28%) (Right). A GFAP⁻/NCAM⁺ neuronal-enriched fraction (P4) is shown for reference. Gates were set with single-stained controls and standard compensation. (N-S) qPCR data of ANLS & Warburg effect signature gene (Group: Saline + SNT, GPI-1&2 + SNT), (each group n=3). Represent fold-change by Saline + Sham. (N) Fold change of *Ldha* in each time point. *P = 0.0310 (Saline + SNT vs Saline + Sham), *P = 0.0108 (Saline + SNT vs GPI-1&2 + SNT). (O) Fold change of *Mct4* in each time point. *P = 0.0312 (Saline + SNT vs Saline + Sham), *P = 0.0118 (Saline + SNT vs GPI-1&2 + SNT). (P) Fold change of *Hk1* in each time point. *P = 0.0243 (Saline + SNT vs Saline + Sham), *P = 0.0102 (Saline + SNT vs GPI-1&2 + SNT). (Q) Fold change of *Pfk1* in each time point. *P = 0.0119(Saline + SNT vs Saline + Sham), **P = 0.0017 (Saline + SNT vs GPI-1&2 + SNT). (R) Fold change of *Pkm* in each time point. (S) Fold change of *Pdha1* in each time point. Data are represented as the mean ± SEM; \**P* < 0.05, **P < 0.01, \*\*\**P* < 0.001, \*\*\*\**P* < 0.0001; Student’s t-value test (H-I and N-S).

Metabolomic clusterings revealed a clear separation of SNT profiles from both the Sham and GPI-treated cohorts, with a substantial contribution from lactate and glutamate (Figures 4B–C). Pathway enrichment analyses showed that metabolic signatures of SNT—including the Warburg effect and glutamate metabolism—were rescued to non-significant levels by glycogenolysis inhibition (Figures 4D–F).

Consistent with effective suppression of glycogen breakdown in both GPI-1 and GPI-2 groups, (Figure 4G, Figure S8), glucose-1-phosphate (G1P) levels were markedly decreased (Figure 4H). Notably, lactate—a hallmark product of the Warburg program—levels were significantly reduced in GPI-treated mice to near-Sham levels (Figure 4I). Additionally, consistent with a glycolytic bias, the lactate/pyruvate ratio was increased in SNT animals and reduced toward Sham levels by both GPI-1 and GPI-2 (Figure S9). At the pathway level, enrichment factors of the Warburg effect and glutamate metabolism—both elevated in SNT—was significantly diminished across multiple statistical metrics in the glycogenolysis-inhibited groups (Figures 4J–K).

To examine gene-level effects of Warburg-like metabolic reprogramming in astrocytes, we isolated ACC astrocytes by FACS following GPI-1/2 treatment and performed transcriptomic profiling (Figure 4L). We sorted astrocytes based on their GFAP-positive versus NCAM-negative status (Figure 4M). Inhibition of glycogenolysis with GPI-1/2 diminished expression of both ANLS and Warburg components: *Ldha*, *Mct4*, *Hk1*, *Pfk1*, and *Pkm*, all elevated in the SNT group, were significantly reduced with GPI treatment (Figures 4M–R). In contrast, *Pdha1*, which channels pyruvate into the TCA cycle, remained unchanged (Figure 4S).

Together, these findings demonstrate that early inhibition of glycogenolysis through PYGB blockage effectively quenches Warburg-type metabolic reprogramming during pain chronification, as evidenced by convergent metabolomic and transcriptomic suppression.

### ACC astrocyte glycogenolysis mediates the chronification of neuropathic pain

Activation of the ANLS in the ACC is essential for pain chronification.^[23–25]^ We therefore asked whether glycogenolysis—the upstream signal that drives Warburg-like metabolic reprogramming— directly contributes to chronic pain and excessive neuronal hyperactivity. To test this, we injected GPI into the contralateral ACC immediately after SNT and assessed outcomes by von Frey behavioral assays and immunohistochemistry (Figure 5A, S10). While GPI treatments did not affect acute nociception, they prevented the nociception in the chronic phase (Figure 5B), indicating that glycogenolysis is required for the persistence, but not the initiation, of pain.

**Figure 5.**
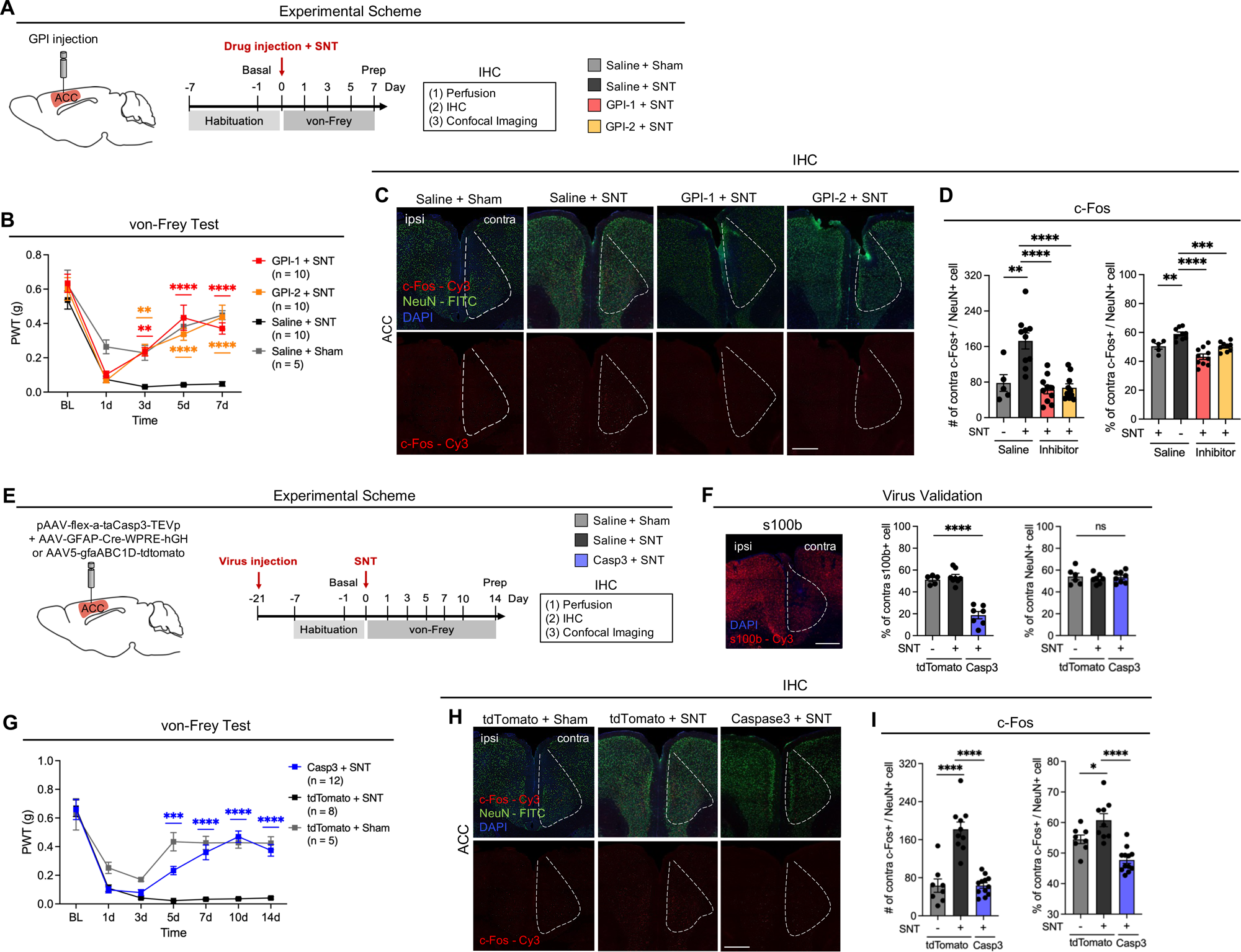
ACC glycogenolysis mediates neuropathic pain chronification. (A) Experimental scheme of NMR and IHC after GPI-1 stereotaxic injection in ACC. (B) von-Frey test of neuropathic pain model. **P = 0.0035 (3 day GPI-1 + SNT vs. Saline + SNT), **P = 0.0018 (3 day GPI-2 + SNT vs. Saline + SNT), ****P < 0.0001 (5 day, 7 day GPI-1 + SNT, GPI-2 + SNT vs. Saline + SNT) (C) Representative confocal images of ACC after IHC (Scale bar, 200 μm). (D) IHC data of c-Fos+/NeuN+ cells in ACC (left): Number of c-Fos+/NeuN+ cells in ACC contra area. **P = 0.0062 (Saline + Sham vs. Saline + SNT), ****P < 0.0001 (GPI-1 + SNT and GPI-2 + SNT vs. Saline + SNT) (right): Ratio of c-Fos+/NeuN+ cells in ACC contra area, compared with ipsi area. **P = 0.0029 (Saline + Sham vs. Saline + SNT), ****P < 0.0001 (GPI-1 + SNT and GPI-2 + SNT vs. Saline + SNT). (E) Experimental Scheme of IHC after AAV-Caspase3 virus injection and SNT surgery. (F) Validation of Caspase3 virus. (Left) Representative confocal images of ACC of anti-S100b staining. (Middle) Ratio of s100b+ cells in ACC contra area, compared with ipsi area. (Shows astrocyte cell density). (Right) Ratio of NeuN+ cells in ACC contra area, compared with ipsi area. (Shows neuron cell density). (G) von-Frey test of neuropathic pain model. \*\*\**P* = 0.0010 (5 day Casp3 + SNT vs. tdTomato + SNT), \*\*\*\**P* < 0.0001 (7 day, 10 day, and 14 day Casp3 + SNT vs. tdTomato + SNT) (H) Representative confocal images of ACC (Scale bar, 200 μm) (I) IHC data of c-Fos+/NeuN+ cells in ACC. tdTomato or Caspase3 injection group analysis (left): Number of c-Fos+/NeuN+ cells in the ACC contralateral area. ****P < 0.0001 (tdTomato + Sham and Casp3 + SNT vs. tdTomato + SNT) (right): Ratio of c-Fos+/NeuN+ cells in ACC contra area, compared with ipsi area. *P = 0.0062 (tdTomato + Sham vs. tdTomato + SNT), ****P < 0.0001 (Casp3 + SNT vs. tdTomato + SNT) Data are represented as the mean ± SEM; \**P* < 0.05, **P < 0.01, \*\*\**P* < 0.001, \*\*\*\**P* < 0.0001; Two-way ANOVA followed by Bonferroni’s multiple comparisons test (B and G) and Student’s t-value test (D, F and I).

Consistent with ACC hyperexcitability as a driver of chronic pain,^[57]^ neuronal c-Fos levels were markedly elevated in SNT mice relative to Sham controls, but significantly reduced by GPI treatment at 7 days (Figures 5C–D). Additional activity markers, including p-CREB and p-p38, followed a similar pattern: elevated in the SNT group yet suppressed by glycogenolysis inhibition (Figures S11A–E). Because the ACC communicates with distributed circuits that underlie pain-related behaviors, we also measured neuronal activity in the nucleus accumbens (NAc) and ventral tegmental area (VTA).^[3,4]^ Both regions exhibited SNT-induced c-Fos upregulation, which was diminished by GPI treatment (Figures S11F–J). Together, these findings suggest that glycogenolysis inhibition selectively reduces pathological neuronal hyperactivity and blocks the chronification of pain, without affecting acute nociceptive responses.

To directly establish the role of astrocytes, we performed astrocyte ablation in the contralateral ACC using a GFAP-promoter–driven Caspase-3 virus (Figure 5E).^[58]^ Expression of GFAP-Casp3 did not alter the number of NeuN-positive neurons, while selectively ablating S100β-positive astrocytes (Figure 5F). Region-specific depletion of astrocytes did not affect acute nociception but prevented the emergence of chronic hypersensitivity (Figures 5G), mirroring the GPI phenotype. Likewise, neuronal c-Fos expression was significantly reduced in the Casp3 group (Figures 5H and 5I).

Finally, we interrogated the contribution of astrocytic lactate shuttling to this process. Inhibition of monocarboxylate transporter (MCTs) with 4-CIN in the contralateral ACC reduced chronic hypersensitivity (Figures S12A–B)^[23,24]^ and suppressed neuronal c-Fos (Figures S12C–D), yielding neuronal activity levels comparable to those seen with GPI treatment (Figure S12E).

Collectively, these results demonstrate that astrocytes drive pain chronification through glycogenolysis-dependent Warburg-like reprogramming, which enhances lactate shuttling and sustains neuronal hyperexcitability (Figure 6). Crucially, this process governs the persistence of pain without affecting acute nociceptive signaling, identifying glycogenolysis as a specific and chemically trackable gatekeeper of chronic pain.

**Figure 6.**
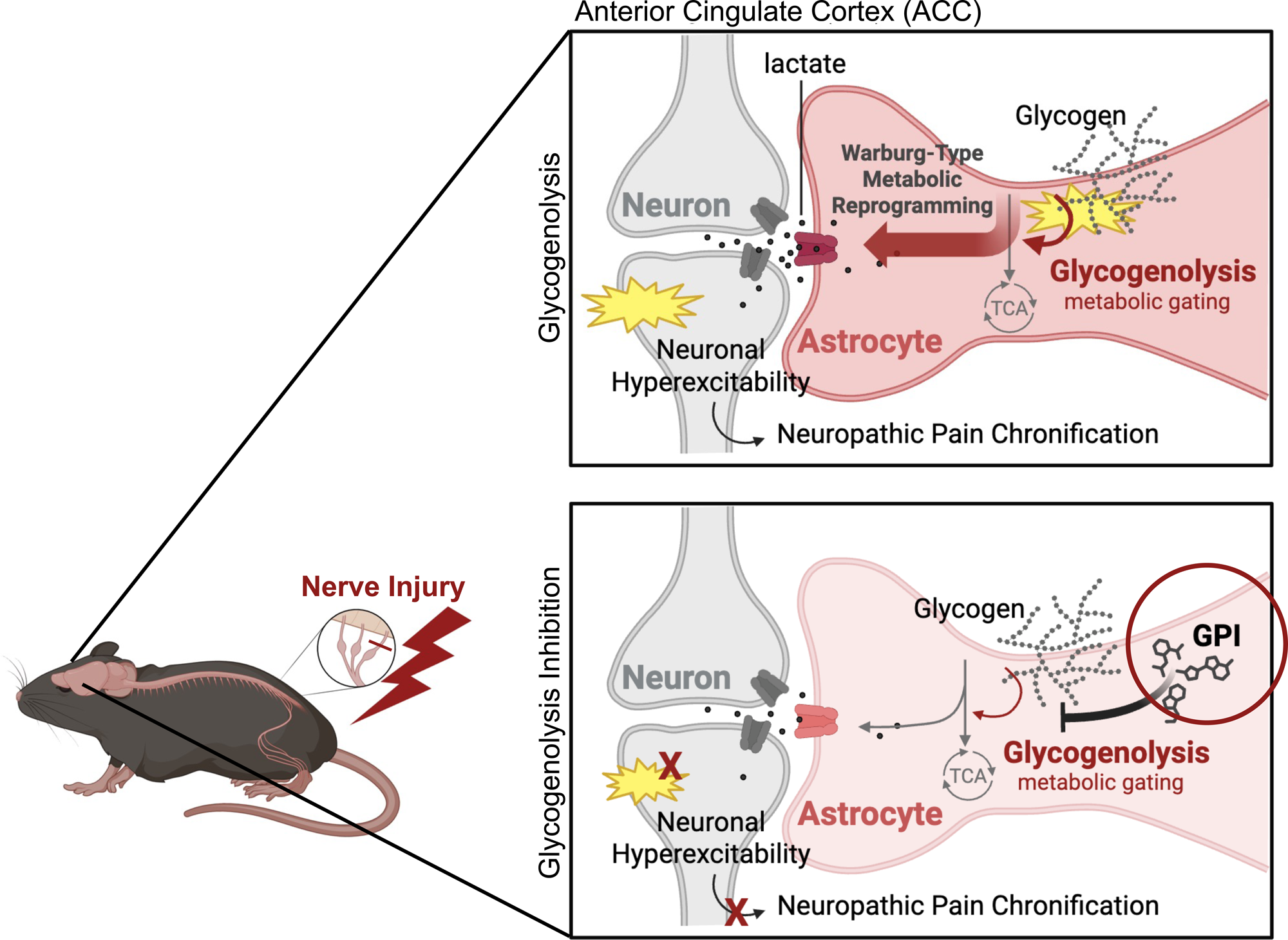
Schematic Illustrations. ACC astrocytic glycogenolysis gates Warburg-like metabolic reprogramming that promotes neuropathic pain chronification.

## DISCUSSION

In chronic neuropathic pain states, astrocytic metabolic adaptations—largely through lactate—have been mainly conceptualized as a fuel that sustains excessive neuronal activity.^[23–25]^ However, we delineate how ACC astrocyte–specific metabolic shifts regulate the chronification of neuropathic pain. By combining time-resolved metabolomics, astrocyte-specific transcriptomics, and targeted chemical perturbations, we uncover a biphasic glycogen program in which early glycogenolysis gates a Warburg-like metabolic reprogramming that supports sustained lactate shuttling and persistent circuit activation. These results reveal cancer-linked, Warburg-like metabolic reprogramming features in CNS astrocytes during chronic pain and nominate glycogen metabolism as a promising entry point for mechanism-based therapeutic strategies in neuropathic pain.

Building on these studies, we extend prior work by moving beyond static end points to provide, to our knowledge, the first time-resolved characterization of how metabolic states in the ACC evolve during the transition from acute nociception to chronic pain. Emerging evidence indicates that diverse astrocytic signaling pathways can rewire metabolic flux, drive transitions to reactive states, and reshape astrocyte outputs. In this light, the time-resolved metabolic reprogramming we describe offers a new vantage point on astrocyte dynamics in chronic neuropathic pain. Rather than a simple, sustained increase in lactate supply, ACC astrocytes in our model exhibit a biphasic metabolic trajectory: an acute bout of glycogenolysis followed by glycogen supercompensation, and a delayed Warburg-like shift as pain becomes chronic. This sequence suggests that astrocytes pass through discrete metabolic states that may encode or reinforce the persistence of neuropathic pain.

Integrating metabolomic and transcriptomic data, we find that glycogenolysis is required for full engagement of the Warburg-like metabolic program in astrocytes in this model, and that blocking this pathway partially normalizes ACC metabolic signatures. From a chemical biology perspective, these results support the view that glycogenolysis functions as an upstream, metabolically defined gate for the Warburg-like state: in our experiments, small-molecule glycogen phosphorylase inhibitors blunt this delayed metabolic reprogramming in astrocytes and concomitantly reduce pain-related circuit activation. Furthermore, ACC astrocytes undergo biphasic regulation of glycogen, and pharmacological inhibition of glycogen breakdown markedly attenuates the development of persistent mechanical hypersensitivity without affecting acute pain sensitization. Thus, glycogen metabolism emerges as an important control node for astrocytic metabolic flux with potential for targeted intervention in neuropathic pain.

During pain chronification, our finding that ACC astrocytes exhibit not only elevated lactate but also broader metabolic remodeling suggests that these adaptations may serve functions beyond simple energetic support. Consistent with this possibility, astrocyte-derived lactate in the ACC may serve as a multifaceted signaling cue that modulates gene expression, neuroimmune interactions, and circuit dynamics, rather than acting solely as a metabolic fuel.^[12,59]^ In hypoxic tumors, elevated lactate production not only supports rapid turnover of metabolic intermediates and redox balance but also mediates histone lactylation, epigenetic remodeling, and inflammatory signaling.^[60]^ Recent studies have further linked astrocyte-intrinsic regulation of glycolysis to the emergence of pro-inflammatory astrocyte states in neuropathic pain.^[61]^ Although these downstream roles remain to be demonstrated directly in chronic pain, our identification of a Warburg-like metabolic state in ACC astrocytes provides a conceptual framework for exploring such non-canonical functions of lactate in neuropathic pain circuits.

The present findings are conceptually aligned with recent spinal cord studies demonstrating that astrocytic glycogen dynamics and neuron–astrocyte metabolic reprogramming contribute to the transition to chronic pain. ^[62–64]^ In these models, distinct glycogen trajectories during pain chronification, together with astrocyte-driven metabolic reprogramming, have been implicated as key determinants of persistent pain states.^[62,63]^ Taken together, these observations support the view that astrocyte-centered metabolic programs represent a more general principle by which glial cells sustain long-lasting activity states in pain pathways.

### Limitations of the study

First, the causal inferences rely on pharmacological inhibition (GPI-1, GPI-2, and 4-CIN), which cannot fully exclude off-target actions or incomplete target engagement; complementary genetic perturbations would further strengthen target validation. Second, our metabolomic and transcriptomic analyses are based on bulk, steady-state measurements and therefore provide system-level snapshots rather than quantitative flux information or single-cell/spatial resolution. Future studies using isotope tracing and single-cell or spatial omics will be important to resolve metabolic heterogeneity among astrocyte subpopulations and their relationships to defined neuronal circuits. Finally, while we highlight glycogenolysis and Warburg-like reprogramming as chemically tractable pathways, the long-term safety and therapeutic feasibility of chronically targeting these metabolic nodes in vivo will need to be evaluated systematically.

## RESOURCE AVAILABILITY

### Lead contact

Further information and requests for resources and reagents should be directed to and will be fulfilled by the lead contact, Prof. Sung Joong Lee or Prof. Seung Bum Park

### Materials availability

All mouse lines and materials used in this study were provided or purchased from the mentioned companies or researchers. This study did not generate any new or unique reagents.

### Data and code availability

All data reported in this paper will be shared by the lead contact upon request.

This paper does not report original code.

All data associated with this study are present in the paper or the supplemental information.

## ACKNOWLEDGMENTS

We are grateful to all members of the Neuron-Glia Network Research Laboratory for their helpful discussions and help. We also thank Kyungchul Noh, assistant professor at the Department of Pharmacology, School of Medicine, Ajou University, Suwon, Republic of Korea, for valuable discussions. This research was supported by the 2024 Seoul National University undergraduate independent research in the Student-Directed Education (SDE) program and the National Research Foundation of Korea (RS-2024-00402116 to S.J.L.; RS-2025-02215169 to S.J.L.; RS-2025-00514527 to S.B.P.).

## AUTHOR CONTRIBUTIONS

J.S.P. designed the research, performed most experiments, analyzed the data, and wrote the first draft of the manuscript. K.H.K. contributed to data interpretation and manuscript revision. H.W.J. crossed RiboTag and GFAP-Cre mice to create astrocyte-specific RiboTag mice and assisted with experimental procedures. S.J.L. and S.B.P. supervised the project and wrote the manuscript.

## METHODS

### Animals

The Institutional Animal Care and Use Committee of Seoul National University approved all animal experiments (Approval No. SNU-250905-4). GFAP-Cre and Rpl22HA/HA (RiboTag) mice were obtained from Jackson Laboratory (GFAP-Cre: B6.Cg-Tg.Gfap-cre.77.6Mvs/2J, Rpl22HA/HA: B6.129-Rpl22tm1.1Psam/J). We generated astrocyte-specific RiboTag mice by crossing floxed RiboTag mice with GFAP-Cre transgenic mice. PCR determined the genotypes of the offspring with the following primers:

*GFAPcre*-Fw: TCC ATA AAG GCC CTG ACA TC;

*GFAPcre*-Rv: TGC GAA CCT CAT CAC TCG T;

*RiboTag*-Fw: GGG AGG CTT GCT GGA TAT G;

*RiboTag*-Rv: TTT CCA GAC ACA GGC TAA GTA CAC.

Except in the RiboTag experiments, all mice used were of the C57BL/6 strain. Male C57BL/6 mice (8–12 weeks of age) were purchased from DooYeol Biotech (Seoul, Korea). All animals were acclimated to standard conditions with a 12-hour light/dark cycle in a specific pathogen-free environment and given access to food and water *ad libitum*. All protocols were performed in accordance with guidelines from the International Association for the Study of Pain.

### Neuropathic pain mouse model

To generate a persistent pain model, a right L5 SNT was performed as previously described. Briefly, animals were anesthetized with isoflurane in an O_2_ carrier (induction 2% and maintenance 1.5%), and a small incision was made to expose the L4 and L5 spinal nerves. The L5 spinal nerve was then transected.

### Reagents

Reagents used in this work include GPI-1 (CP-316819, CAS 186392-43-8) from TOCRIS, GPI-2 (361515-1MG, CAS 648926-15-2), 4-CIN (α-cyano-4-hydroxycinnamic acid, CAS 28166-41-8) from Sigma-Aldrich. Each chemical was prepared as a 100× stock in DMSO and diluted as required for use.

### Pain behavior test (von Frey test)

Mechanical sensitivity of the right hind paw was assessed using a calibrated series of von Frey hairs (0.02–6 g, Stoelting, Wood Dale, IL, USA) following the up-down method.^[65,66]^ Tests were performed after at least three habituations, each at 24-hour intervals. Assessments were made 1 day before surgery for baseline, and 1, 3, 5, and 7 days after SNT. Rapid paw withdrawal, licking, and flinching were interpreted as signs of pain. All behavioral tests were performed in a blinded manner to the conditions.

### Stereotaxic injection

For stereotaxic drug injection, each C57BL/6 mouse received a unilateral injection of 1 μl of GPI-1 at 500 nM (CP-316819, CAS 186392-43-8; TOCRIS, Minneapolis, MN, USA), 1 μl of GPI-2 at 500 nM (361515-1MG, CAS 648926-15-2; Sigma-Aldrich, St. Louis, MO, USA), or 1 μl of 4-CIN at 0.5 mM (α-cyano-4-hydroxycinnamic acid, CAS 28166-41-8; Sigma-Aldrich) in the ACC using the following coordinates: AP, −1.0 mm; ML, 0.4 mm; DV, −1.5 mm from the bregma. Drug injected immediately after SNT surgery.

For stereotaxic virus injection, each C57BL/6 mouse received a unilateral injection of 1 μl of pAAV-flex-a-taCasp3-TEVp, AAV-GFAP-Cre-WPRE-hGH, or AAV5-gfaABC1D-tdTomato (~1 × 10^13^ gene copies (GC)/ml) in the ACC using the following coordinates: AP, −0.9 mm; ML, 0.4 mm; DV, −1.5 mm from the bregma. SNT surgery was performed 4 weeks after the virus injections. The injection syringe (Hamilton, Reno, NV, USA) delivered GPI or AAV at a constant volume of 0.1 μl/min using a syringe pump (Stoelting, Wood Dale, IL, USA).

### Immunohistochemistry (IHC)

Mice were transcardially perfused with ice-cold 0.1 M phosphate-buffered saline (PBS; pH 7.4) until all blood was removed, followed by perfusion with ice-cold 4% paraformaldehyde in 0.1 M PBS. Whole brains were post-fixed in 4% paraformaldehyde in 0.1 M PBS overnight at 4°C and cryoprotected with 30% sucrose for 3 days. Coronal 60-μm-thick sections were incubated in cryoprotectant at −20°C until immunohistochemical staining was performed. The sections were incubated for 1 h at room temperature in a blocking solution containing 5% normal goat serum (Jackson ImmunoResearch, Bar Harbor, ME, USA), 2% BSA (Sigma-Aldrich), and 0.1% Triton X-100 (Sigma-Aldrich).

Subsequently, the sections were incubated in the blocking solution with mouse anti-NeuN (MAB377B, 1:1000; Millipore, Billerica, MA, USA), rabbit anti-S100b (ab52642, 1:500; Abcam, Cambridge, MA, USA), mouse anti-phospho-CREB (#9198 87G3, 1:1000; Cell Signaling Technology, Danvers, MA, USA) rabbit anti-p-p38 MAPK (# 9211S, 1:1000; Cell Signaling Technology), or rabbit anti-c-Fos (#2250 9F6, 1:1000; Cell Signaling Technology) antibodies overnight at 4°C. After being washed with 0.1 M PBS containing 0.1% Triton X-100, the sections were incubated in blocking solution for 1 h with FITC-, Cy3- or Cy5-conjugated secondary antibodies (1:200, Jackson ImmunoResearch) at room temperature, washed three times, and then mounted on gelatin-coated glass slides using Vectashield (Vector Laboratories, Inc., Burlingame, CA, USA). Fluorescent images of the mounted sections were obtained using a confocal microscope (LSM800; Carl Zeiss, Jena, Germany).

### Glycogen assay

#### Sample preparation

For brain sections, the animals were killed under isoflurane, and the brain was quickly removed from the skull and immediately frozen with dry ice. ACC sections were collected based on measurements from the Allen Brain Atlas, and the samples were snap-frozen and stored at −80°C until further processing.

#### Glycogen colorimetric assay

Tissue samples were homogenized on ice in 200 µL of ddH₂O using a Dounce homogenizer with 10–15 passes. Homogenates were boiled for 10 min to inactivate enzymes and subsequently centrifuged at 18,000 × g for 10 min at 4°C to remove insoluble material. The resulting supernatant was collected and used for the glycogen analysis. To perform the assay, 2–50 µL of the tissue supernatant was added to a 96-well plate, and the volume was adjusted to 50 µL per well with glycogen hydrolysis buffer provided in a glycogen colorimetric/fluorometric assay kit (ab65620, Abcam). For glycogen detection, 2 µL of hydrolysis enzyme mix was added to the wells designated for glycogen hydrolysis, and the background control wells received no enzyme. The samples were incubated at room temperature for 30 min to allow the hydrolysis of glycogen to glucose.

Following hydrolysis, 50 µL of a reaction mix containing 46 µL of development buffer, 2 µL of development enzyme mix, and 2 µL of OxiRed probe was added to all wells. The plates were then incubated in the dark at room temperature for 30 min. Absorbance was measured at 570 nm using a microplate reader to quantify the glycogen content. A standard curve was generated using glycogen standards (0–2 µg/well) prepared according to the manufacturer’s instructions to determine the sample glycogen concentrations. Background absorbance from control wells was subtracted from the sample wells to account for any non-glycogen-derived signal. The glycogen concentration in each sample was normalized to the initial sample volume and adjusted according to the dilution factors.

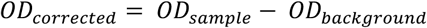

where OD sample is the measured OD value for the sample well, and OD background is the measured OD value for the sample background control well (except in the hydrolysis mix, where only the background glucose constant was measured).

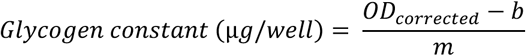

where m is the slope of the standard curve (glycogen constant), b is the intercept of the standard curve, and OD corrected is the corrected OD value for the sample well.

### Western blotting

For Western blotting, ACC tissues were homogenized in ice-cold RIPA buffer (50 mM Tris-HCl, pH 7.5, 150 mM NaCl, 1% NP-40, 0.5% sodium deoxycholate, 0.1% SDS) supplemented with 1 mM PMSF and a phosphatase inhibitor cocktail (Sigma-Aldrich, P5726). The homogenates were incubated on ice for 30 min and then centrifuged at 13,000 rpm for 15 min at 4°C. The supernatants were then collected. Protein concentration was measured using a BCA assay (Pierce, 23225). For each sample, 20 µg was mixed with 5× SDS sample buffer, boiled for 5 min at 95°C, and then resolved on a 10% SDS-PAGE gel. Proteins were transferred to nitrocellulose membranes (LC2001; Invitrogen, Carlsbad, CA, USA) at 100 V for 1 h, blocked in 5% milk/TBST for 1 h, and probed overnight at 4°C with mouse anti-HA (ab9110, 1:2000; Abcam) and mouse anti-β-actin (A2228, 1:5000; Sigma-Aldrich) in 2.5% milk/TBST. After three TBST washes, the membranes were incubated with HRP-conjugated goat anti-mouse IgG (1:3000 in 2.5% milk/TBST) for 1 h at room temperature. Blots were developed using SuperSignal™ West Pico PLUS (Thermo Fisher, Waltham, MA, USA), and images were captured using a Fusion FX6.0 system. Band intensities were quantified in ImageJ and normalized to β-actin.

### qPCR

The real-time RT-PCR (qPCR) experiments were performed using a StepOnePlus real-time PCR system (Applied Biosystems, Foster City, CA, USA) following the 2−ΔΔCt method. Total RNA from contralateral ACC tissue was extracted using TRIzol (Invitrogen) and reverse transcribed using TOPscript RT DryMIX (Enzynomics, Cat # RT200, Daejeon, Korea). All the ΔCt values were normalized to the corresponding GAPDH values, and represent fold change induction.

The following qPCR primers were used:

*Ppp1r3c*-Fw: GGT GAC TCA TCT TTC TGC CAC A;

*Ppp1r3c*-Rv: CAA GAC AAA ATT AGG CAC GAG A;

*Gys1*-Fw: ATC TAC ACT GTG CTG CAG ACG;

*Gys1*-Rv: CCC TTG CTG TTC ATG GAA TCC;

*Phka2*-Fw: TGG ATG CCA CCT CTC TCT TC;

*Phka2*-Rv: TAT CTC CAC GCT CCC ACA TC;

*Pygb*-Fw: CAG CAG CAT TAC TAT GAG CGG;

*Pygb*-Rv: CCA AGT CCA ACC CCA ACT GA;

*Hif-1a*-Fw: GAT CCT TGA TGC TTG CTG GG;

*Hif-1a*-Rv: CTG TCC CCA ATG TCC AGA GT;

*Ldha*-Fw: AAA GAG GAC TAA GGG GTG GC;

*Ldha*-Rv: CTG CAG GAA ACA ACC ACT CC;

*Ldhb*-Fw: AAA GGC TAC ACC AAC TGG GC;

*Ldhb*-Rv: GCC GTA CAT TCC CTT CAC CA;

*Mct4*-Fw: CAT TCC CAG GGA CGC AAA GAG;

*Mct4*-Rv: GAC ACG GCT TGG ATC TCC TC;

*HK1*-Fw: CCA TCC CTC TTT GAC ACC CT;

*HK1*-Rv: ACT CAG ACT AAA GTG GCC CC;

*Pfk1*-Fw: CAG AAA GCC CAC ACT CAA CC;

*Pfk1*-Rv: ACA GAA GAC CTT GGC CTA CC;

*Pkm*-Fw: CTG GGT GGG AGA AAT GGA GT;

*Pkm*-Rv: TCA GAA GCC CAG AGA ACC AG;

*Pdha1*-Fw: GAT GCC GTG CTG ATT TAG GG;

*Pdha1*-Rv: CGT CCT AGA AAT GGC AGC AC.

### pH-analysis

Following dissection of the anterior cingulate cortex (ACC), tissue was homogenized in ultrapure distilled water (pH 7.0) using a cell homogenizer. Homogenates were centrifuged to pellet cellular debris, and the pH of the resulting supernatant was measured with a FiveEasy pH meter (F20-Std-Kit; Mettler-Toledo, Greifensee, Switzerland). For each specimen, pH was recorded in triplicate, and the mean value was used for analysis.

### FACS Cell Sorting

Anterior cingulate cortex (ACC) tissue was prepared in FACS buffer (0.1% BSA in PBS) and incubated for 20 min in buffer containing collagenase A (0.5 mg/mL). Following gentle trituration, the suspension was passed through a nylon cell strainer to obtain single cells. Cells were then sequentially labeled in FACS buffer at 4 °C for 20 min each with primary antibodies—rabbit anti-GFAP (Dako, Glostrup, Denmark) and mouse anti-NCAM (MilliporeSigma, Burlington, MA, USA)—followed by secondary antibodies—donkey anti-rabbit Cy5 and donkey anti-mouse FITC (Jackson ImmunoResearch, West Grove, PA, USA). Labeled cells were analyzed and sorted on a FACSAria Fusion (BD Biosciences, San Jose, CA, USA), gating on live, singlet events to isolate GFAP⁺/NCAM⁻ cells. Sorted cells were subsequently used for quantitative PCR (qPCR).^[67]^

### RNA-seq data analysis

Raw RNA-seq data were processed in Python to analyze relative gene expression between the experimental and control groups. Raw counts were compared against the sham group and normalized for library size using DESeq2’s size-factor adjustment. A negative binomial model was then fitted to calculate the log₂ fold change and the associated *p-*value for each gene. Genes were ranked in descending order by their log₂FC/standard error, and a GSEA was performed against the Hallmark and Gene Ontology gene sets. Normalized enrichment scores and false discovery rates were computed, and enrichment plots were generated in GraphPad Prism. For each analysis, we selected gene sets corresponding to specific metabolic pathways to assess their degree of enrichment.

### GC-MS

Tissue samples for the GC-MS analysis were prepared from the ACC using the same dissection and freezing protocol as for the glycogen assay. For each contralateral ACC specimen, 400 µL of tissue supernatant was collected and subjected to GC-MS. Derivatization was performed using trimethylsilylation, and 1 µL of each derivatized sample was injected into a Thermo Scientific ISQ LT GC-MS system (Thermo Scientific, Waltham, MA, USA).^[27]^ Chromatographic separation and mass detection were performed under standard operating conditions. Raw data were processed using Thermo Xcalibur Quan Browser. Metabolites were identified by matching each chromatographic peak to reference spectra, and peak areas were integrated against the baseline to obtain relative signal intensities. All metabolite abundances were then normalized to the sham controls and expressed as fold changes for downstream analysis.

### NMR

Tissue for the NMR analyses of the ACC was prepared using the same dissection and freezing protocol as for the glycogen assay. For each contralateral ACC sample, 500 µL of the tissue supernatant was collected and analyzed by NMR.^[68,69]^ Spectra were acquired on a Bruker AVANCE III HD 600 MHz high-resolution NMR spectrometer (Bruker BioSpin, Rheinstetten, Germany). Individual resonances were assigned using Chenomx NMR Suite Profiler, and metabolite concentrations were determined by integrating each peak relative to the subtraction baseline.

### Metabolomics analysis

Absolute and relative metabolite concentrations obtained by GC-MS and NMR, including fold-change values normalized to the sham controls, were subjected to a comprehensive metabolomics analysis. For each metabolite, both the absolute abundance and the fold change relative to the sham and saline + SNT groups were calculated, and statistical significance was assessed. To identify enriched metabolic pathways within each experimental group, we performed both an over-representation analysis and a quantitative enrichment analysis. All pathway enrichment analyses were conducted using MetaboAnalyst v6.0. (http://www.metaboanalyst.ca).^[36,37]^

### In silico analysis

#### PYGB interaction analysis

The crystal structure of PYGB (PDB ID: 5IKP) was downloaded from the RCSB Protein Data Bank (https://www.rcsb.org/structure/5IKP) and used as the starting model for all subsequent structural analyses. Ligand coordinates were generated in silico from the canonical SMILES string. Hydrogens were added, and the geometry was energy-minimized to convergence; the lowest-energy conformer was exported in PDB format. Protein coordinates (PDB) were parsed with a structural biology toolkit to extract atom and residue-level information. Ligand residues were recognized by residue name, and neighboring amino acids within a predefined distance cutoff were designated as pocket residues. These pocket residues were recorded for subsequent validation and docking calculations to characterize protein–ligand interactions. Protein–ligand docking simulations were performed using GalaxyDockWeb and CBDock2.^[44,47]^

#### Glycogen phosphorylase inhibitor selection

We evaluated six commonly used glycogen phosphorylase inhibitors for their predicted binding affinity to PYGB. Canonical SMILES strings for each compound were used to generate 3D ligand structures, which were then submitted to CBDock2 (default settings).^[44–46]^ For each ligand–protein pair, docking scores were reported as predicted binding free energies (ΔG, kcal/mol). For PYGB, ΔG values were calculated both for a predefined reference binding site and across all binding cavities detected by the algorithm. Representative docking poses in the selected pocket are shown together with the corresponding contact residues for all six compounds.

Among these, two compounds (GPI-1 and GPI-2) exhibited the most favorable (lowest) predicted ΔG values and well-defined interaction networks within the reference pocket, and were therefore selected as lead PYGB inhibitors. The binding poses of GPI-1 and GPI-2 were visualized using UCSF ChimeraX (Resource for Biocomputing, Visualization, and Informatics, University of California, San Francisco; RRID:SCR_015872) (Fig. 3M).

#### Enzyme isoform validation analysis

To assess enzyme isoform selectivity, the crystal structures of PYGL (PDB ID: 2QLL) and PYGM (PDB ID: 1Z8D) were downloaded from the RCSB Protein Data Bank (https://www.rcsb.org/structure/2QLL and https://www.rcsb.org/structure/1Z8D) and used as starting models for structural analysis; for PYGB, the same structure as in the PYGB interaction analysis was used (PDB ID: 5IKP). Docking of GPI-1 and GPI-2 to PYGB, PYGL, and PYGM was performed in CBDock2 with the same parameters as in the primary PYGB analysis. For each isoform, predicted binding free energies (ΔG, kcal/mol) were computed for the predefined reference pocket as well as for all identified binding sites, allowing comparison of the relative affinity of GPI-1 and GPI-2 across glycogen phosphorylase isoforms.

#### Off-target validation analysis (Target Screening)

Two small molecules (GPI-1 and GPI-2) were profiled using GalaxySagittarius-AF (default settings).^[47–49]^ The platform selects a binding pocket per UniProt target and returns (i) a Predock score (unitless; higher indicates greater ligand–pocket compatibility) and (ii) a GalaxyDock BP2 docking energy (scoring-function value; more negative is better). Ligands were standardized by enumerating relevant protomer/tautomer states at pH 7.4 ± 0.5 and forwarding the dominant state.

For each ligand’s table, scores were normalized within-sheet to place components on a common 0–1 scale:

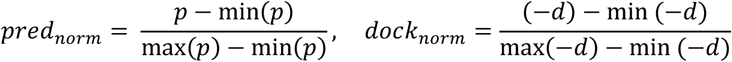

Where p is Predock and d is the docking energy (sign-flipped so that larger is better). The platform’s composite score was then reconstructed by ordinary least squares (OLS) per sheet as:

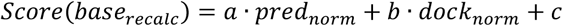

With coefficients (a,b,c) fit to the sheet data (R^2^ ≈ 1 for both ligands).

Because both compounds are glycogen phosphorylase (GP) inhibitors, we applied a small heuristic class adjustment based on target class (UniProt mapping). The Final priority score used for ranking and plotting was:

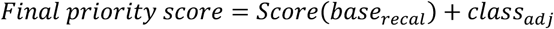

With additive adjustments: GP +0.05, kinase −0.08, serine protease −0.10, metalloprotease −0.10, HSP90 −0.05, carbonic anhydrase −0.06, nuclear receptor −0.04, albumin −0.06, unknown 0. GP isoforms were defined as PYGL (P06737), PYGM (P11217), and PYGB (P11216).

Targets were rank-ordered by the Final priority score (y-axis; Figures S5D–E). GP isoforms were highlighted in red, and insets report raw Predock score and Docking energy (GalaxyDock BP2; Figures S5D-E) for the top GP hits.

### Quantification and statistical analysis

The data were analyzed in GraphPad software. Student’s *t-*test was used for comparisons between two groups. For multiple group comparisons, two-way analysis of variance (ANOVA) was conducted, followed by Bonferroni’s post hoc test. All data are expressed as the mean ± standard error of the mean (SEM), and statistical significance was defined as a p-value < 0.05.

## REFERENCES

1. Bliss TVP, Collingridge GL, Kaang B-K, Zhuo M. (2016). Synaptic plasticity in the anterior cingulate cortex in acute and chronic pain. Nat Rev Neurosci. 17, 485–496.

2. Zhuo M. (2008). Cortical excitation and chronic pain. Trends Neurosci. 31, 199–207.

3. Song Q, Wei A, Xu H, et al. (2024). An ACC–VTA–ACC positive-feedback loop mediates the persistence of neuropathic pain and emotional consequences. Nat Neurosci. 27, 272–285.

4. Gao SH, Shen LL, Wen HZ, Zhao YD, Chen PH, Ruan HZ. (2020). The projections from the anterior cingulate cortex to the nucleus accumbens and ventral tegmental area contribute to neuropathic pain-evoked aversion in rats. Neurobiol Dis. 140, 104862.

5. Nagai J, Yu X, Papouin T, et al. (2021). Behaviorally consequential astrocytic regulation of neural circuits. Neuron. 109, 576–596.

6. Bolaños JP, Magistretti PJ. (2025). The neuron–astrocyte metabolic unit as a cornerstone of brain energy metabolism in health and disease. Nat Metab. (2025).

7. Zhang Y-M, Qi Y-B, Gao Y-N, Chen W-G, Zhou T, Zang Y, Li J. (2023). Astrocyte metabolism and signaling pathways in the CNS. Front Neurosci. 17, 1217451.

8. Bonvento G, Bolaños JP. (2021). Astrocyte-neuron metabolic cooperation shapes brain activity. Cell Metab. 33, 1546–1564.

9. Theparambil SM, Kopach O, Braga A, et al. (2024). Adenosine signalling to astrocytes coordinates brain metabolism and function. Nature. 632, 139–146.

10. Schreiber SL, Kotz JD, Li M, et al. (2015). Advancing biological understanding and therapeutics discovery with small-molecule probes. Cell. 161, 1252–1265.

11. Gerry CJ, Schreiber SL. (2018). Chemical probes and drug leads from advances in synthetic planning and methodology. Nat Rev Drug Discov. 17, 333–352.

12. Magistretti PJ, Allaman I. (2018). Lactate in the brain: from metabolic end-product to signalling molecule. Nat Rev Neurosci. 19, 235–249.

13. Kim Y, Dube SE, Park CB. (2025). Brain energy homeostasis: the evolution of the astrocyte–neuron lactate shuttle hypothesis. Korean J Physiol Pharmacol. 29, 1–8.

14. Suzuki A, Stern SA, Bozdagi O, et al. (2011). Astrocyte-neuron lactate transport is required for long-term memory formation. Cell. 144, 810–823.

15. Wang J, Tu J, Cao B, et al. (2017). Astrocytic L-lactate signaling facilitates amygdala–anterior cingulate cortex synchrony and decision making in rats. Cell Rep. 21, 2407–2418.

16. Clasadonte J, Scemes E, Wang Z, Boison D, Haydon PG. (2017). Connexin 43-mediated astroglial metabolic networks contribute to the regulation of the sleep–wake cycle. Neuron. 95, 1365–1380.e5.

17. Descalzi G, Gao V, Steinman MQ, Suzuki A, Alberini CM. (2019). Lactate from astrocytes fuels learning-induced mRNA translation in excitatory and inhibitory neurons. Commun Biol. 2, 247.

18. Natsubori A, Hirai S, Kwon S, et al. (2023). Serotonergic neurons control cortical neuronal intracellular energy dynamics by modulating astrocyte-neuron lactate shuttle. iScience. 26, 105830.

19. Akter M, Hasan M, Ramkrishnan AS, et al. (2023). Astrocyte and L-lactate in the anterior cingulate cortex modulate schema memory and neuronal mitochondrial biogenesis. eLife. 12, e85751.

20. Cauli B, Dusart I, Li D. (2023). Lactate as a determinant of neuronal excitability, neuroenergetics and beyond. Neurobiol Dis. 184, 106207.

21. Braga A, Chiacchiaretta M, Pellerin L, Kong D, Haydon PG. (2024). Astrocytic metabolic control of orexinergic activity in the lateral hypothalamus regulates sleep and wake architecture. Nat Commun. 15, 5979.

22. Fernández-Moncada I, Lavanco G, Fundazuri UB, et al. (2024). A lactate-dependent shift of glycolysis mediates synaptic and cognitive processes in male mice. Nat Commun. 15, 6842.

23. Wang Y, Peng Y, Zhang C, Zhou X. (2021). Astrocyte-neuron lactate transport in the anterior cingulate cortex contributes to the occurrence of long-lasting inflammatory pain in male mice. Neurosci Lett. 764, 136205.

24. Iqbal Z, Liu S, Lei Z, Ramkrishnan AS, Akter M, Li Y. (2023). Astrocyte L-lactate signaling in the anterior cingulate cortex regulates visceral pain aversive memory in rats. Cells. 12, 26.

25. Reid P, Scherer K, Halasz D, et al. (2025). Astrocyte neuronal metabolic coupling in the anterior cingulate cortex of mice with inflammatory pain. Brain Behav Immun. 125, 212–225.

26. Alberini CM, Cruz E, Descalzi G, Bessières B, Gao V. (2018). Astrocyte glycogen and lactate: New insights into learning and memory mechanisms. Glia. 66, 1244–1262.

27. Gibbs ME, Anderson DG, Hertz L. (2006). Inhibition of glycogenolysis in astrocytes interrupts memory consolidation in young chickens. Glia. 54, 214–222.

28. Bak LK, Walls AB, Schousboe A, Waagepetersen HS. (2018). Astrocytic glycogen metabolism in the healthy and diseased brain. J Biol Chem. 293, 7108–7116.

29. Vander Heiden MG, Cantley LC, Thompson CB. (2009). Understanding the Warburg effect: the metabolic requirements of cell proliferation. Science. 324, 1029–1033.

30. Pavlova NN, Thompson CB. (2016). The emerging hallmarks of cancer metabolism. Cell Metab. 23, 27–47.

31. DeBerardinis RJ, Chandel NS. (2016). Fundamentals of cancer metabolism. Sci Adv. 2, e1600200.

32. Lee J, Noh K, Lee S, et al. (2025). Ganglioside GT1b prevents selective spinal synapse removal following peripheral nerve injury. EMBO Rep. 26, 2994–3023.

33. Lee J, Lee G, Ko G, Lee SJ. (2023). Nerve injury-induced gut dysbiosis contributes to spinal cord TNF-α expression and nociceptive sensitization. Brain Behav Immun. 110, 155–161.

34. Lim H, Lee J, You B, et al. (2020). GT1b functions as a novel endogenous agonist of toll-like receptor 2 inducing neuropathic pain. EMBO J. 39, e102214.

35. Chan ECY, Pasikanti KK, Nicholson JK. (2011). Global urinary metabolic profiling procedures using gas chromatography–mass spectrometry. Nat Protoc. 6, 1483–1499.

36. Pang Z, Lu Y, Zhou G, et al. (2024). MetaboAnalyst 6.0: towards a unified platform for metabolomics data processing, analysis and interpretation. Nucleic Acids Res. 52, W398–W406.

37. Ewald JD, Zhou G, Lu Y, et al. (2024). Web-based multi-omics integration using the Analyst software suite. Nat Protoc. 19, 1467–1497.

38. Zhang Y, Jiang S, Liao F, et al. (2022). A transcriptomic analysis of neuropathic pain in the anterior cingulate cortex after nerve injury. Bioengineered. 13, 2058–2075.

39. Zhang Y, Chen K, Sloan SA, et al. (2014). An RNA-sequencing transcriptome and splicing database of glia, neurons, and vascular cells of the cerebral cortex. J Neurosci. 34, 11929–11947.

40. Canada SE, Weaver SA, Sharpe SN, Pederson BA. (2011). Brain glycogen supercompensation in the mouse after recovery from insulin-induced hypoglycemia. J Neurosci Res. 89, 585–591.

41. Jakobsen E, Bak LK, Walls AB, Reuschlein AK, Schousboe A, Waagepetersen HS. (2017). Glycogen shunt activity and glycolytic supercompensation in astrocytes may be distinctly mediated via the muscle form of glycogen phosphorylase. Neurochem Res. 42, 2490–2494.

42. Xie H, Song J, Godfrey J, et al. (2021). Glycogen metabolism is dispensable for tumour progression in clear cell renal cell carcinoma. Nat Metab. 3, 327–336.

43. Ibrahim MMH, Bheemanapally K, Alhamami HN, Briski KP. (2020). Effects of intracerebroventricular glycogen phosphorylase inhibitor CP-316,819 infusion on hypothalamic glycogen content and metabolic neuron AMPK activity and neurotransmitter expression in male rat. J Mol Neurosci. 70, 647–658.

44. Liu Y, Yang X, Gan J, Liu D, Cao Y. (2022). CB-Dock2: improved protein–ligand blind docking by integrating cavity detection, docking and homologous template fitting. Nucleic Acids Res. 50, W159–W164.

45. Yang X, Liu Y, Gan J, Xiao Z-X, Cao Y. (2022). FitDock: protein–ligand docking by template fitting. Brief Bioinform. 23, bbac087.

46. Gan J-H, Liu J-X, Liu Y, Chen S-W, Dai W-T, Xiao Z-X, Cao Y. (2023). DrugRep: an automatic virtual screening server for drug repurposing. Acta Pharmacol Sin. 44, 888–896.

47. Ko J, Park H, Heo L, Seok C. (2012). GalaxyWEB server for protein structure prediction and refinement. Nucleic Acids Res. 40, W294–W297.

48. Kwon S, Jung N, Yang J, Seok C. (2024). GalaxySagittarius-AF: Predicting targets for drug-like compounds in the extended human 3D proteome. J Mol Biol. 436, 168617.

49. Yang J, Kwon S, Bae S-H, Park KM, Yoon C, Lee J-H, Seok C. (2020). GalaxySagittarius: Structure-and similarity-based prediction of protein targets for druglike compounds. J Chem Inf Model. 60, 3246–3254.

50. Kuleshov MV, Jones MR, Rouillard AD, et al. (2016). Enrichr: a comprehensive gene set enrichment analysis web server 2016 update. Nucleic Acids Res. 44, W90–W97.

51. Xie Z, Bailey A, Kuleshov MV, et al. (2021). Gene set knowledge discovery with Enrichr. Curr Protoc. 1, e90.

52. Favaro E, Bensaad K, Chong MG, et al. (2012). Glucose utilization via glycogen phosphorylase sustains proliferation and prevents premature senescence in cancer cells. Cell Metab. 16, 751–764.

53. Zois CE, Harris AL. (2016). Glycogen metabolism has a key role in the cancer microenvironment and provides new targets for cancer therapy. J Mol Med. 94, 137–154.

54. Shulman RG, Rothman DL. (2017). The glycogen shunt maintains glycolytic homeostasis and the Warburg effect in cancer. Trends Cancer. 3, 761–767.

55. Curtis M, Kenny HA, Ashcroft B, et al. (2019). Fibroblasts mobilize tumor cell glycogen to promote proliferation and metastasis. Cell Metab. 29, 141–155.e9.

56. Liu Q, Li J, Zhang W, et al. (2021). Glycogen accumulation and phase separation drives liver tumor initiation. Cell. 184, 5559–5576.e19.

57. Wei N, Guo Z, Qiu M, et al. (2024). Astrocyte activation in the ACC contributes to comorbid anxiety in chronic inflammatory pain and involves in the excitation–inhibition imbalance. Mol Neurobiol. 61, 6934–6949.

58. Chan KY, Jang MJ, Yoo BB, et al. (2017). Engineered AAVs for efficient noninvasive gene delivery to the central and peripheral nervous systems. Nat Neurosci. 20, 1172–1179.

59. Xiong XY, Tang Y, Yang QW. (2022). Metabolic changes favor the activity and heterogeneity of reactive astrocytes. Trends Endocrinol Metab. 33, 390–400.

60. Li X, Yang Y, Zhang B, et al. (2022). Lactate metabolism in human health and disease. Signal Transduct Target Ther. 7, 305.

61. Chen Y, Liao Y, Zhu H, et al. (2025). Sox9 regulation of hexokinase 1 controls neuroinflammatory astrocyte subtypes in a rat model of neuropathic pain. Nat Commun. 16, 10249.

62. Díaz-García CM. (2024). Glycogen from spinal astrocytes dials up the pain. Nat Metab. 6, 384–386.

63. Marty-Lombardi S, Lu S, Ambroziak W, et al. (2024). Neuron-astrocyte metabolic coupling facilitates spinal plasticity and maintenance of inflammatory pain. Nat Metab. 6, 494–513.

64. Mabou Tagne A, Fotio Y, Lee HL, et al. (2025). Metabolic reprogramming in the spinal cord drives the transition to pain chronicity. Cell Rep. 44, 116261.

65. Tanga FY, Nutile-McMenemy N, DeLeo JA. (2005). The CNS role of toll-like receptor 4 in innate neuroimmunity and painful neuropathy. Proc Natl Acad Sci U S A. 102, 5856–5861.

66. Chaplan SR, Bach FW, Pogrel JW, Chung JM, Yaksh TL. (1994). Quantitative assessment of tactile allodynia in the rat paw. J Neurosci Methods. 53, 55–63.

67. Sergent-Tanguy S, Chagneau C, Neveu I, Naveilhan P. (2003). Fluorescent activated cell sorting (FACS): a rapid and reliable method to estimate the number of neurons in a mixed population. J Neurosci Methods. 129, 73–79.

68. Crook AA, Powers R. (2020). Quantitative NMR-based biomedical metabolomics: current status and applications. Molecules. 25, 5128.

69. Pudełko-Malik N, Drulis-Fajdasz D, Pruss Ł, et al. (2024). A single dose of glycogen phosphorylase inhibitor improves cognitive functions of aged mice and affects the concentrations of metabolites in the brain. Sci Rep. 14, 24123.

