## Supplementary Information for "Astrocytic glycogenolysis gates Warburg-like metabolic reprogramming that promotes neuropathic pain chronification"

| Contents | Page |
|---|---|
| Key Resources Details | 2 |
| Methods and Materials | 3 |
| Supplementary Figures | 6 |
| Methods Validations | 21 |

### Key Resources Detail

| REAGENT OR RESOURCE | SOURCE | IDENTIFIER |
|---|---|---|
| Antibodies | | |
| Mouse anti-NeuN (clone A60) | Millipore Sigma | Cat# MAB377B |
| Rabbit anti-S100β | Abcam | Cat# ab52642 |
| Rabbit anti-phospho-CREB | Cell Signaling Technology | Cat# 9198 |
| Rabbit anti-phospho-p38 MAPK | Cell Signaling Technology | Cat# 9211 |
| Rabbit anti-c-Fos (9F6) | Cell Signaling Technology | Cat# 2250 |
| Rabbit anti-GFAP | DAKO | Cat# Z0334 |
| Mouse anti-PSA-NCAM | Millipore Sigma | Cat# MAB5234 |
| Rabbit anti-HA tag | Abcam | Cat# ab9110 |
| Mouse anti-β-Actin | Sigma-Aldrich | Cat# A2228 |
| FITC / Cy3 / Cy5-conjugated secondary antibodies | Jackson ImmunoResearch | Cat# 115-095-062, 111-165-008 |
| HRP-conjugated goat anti-mouse IgG | Jackson ImmunoResearch | Cat# 115-035-003 |
| Bacterial and virus strains | | |
| AAV5-flex-taCasp3-TEVp | Addgene | Cat# 45580-AAV5 |
| AAV5-GFAP-Cre-WPRE-hGH | Addgene | Cat# 105550-AAV5 |
| AAV5-gfaABC1D-tdTomato | Addgene | Cat# 44332-AAV5 |
| Biological samples | | |
| Mouse anterior cingulate cortex (ACC) tissue | This paper | N/A |
| Chemicals, peptides, and recombinant proteins | | |
| CP-316819 (GPI-1) | Tocris | Cat# 3542 |
| Glycogen phosphorylase inhibitor (GPI-2) | Sigma-Aldrich | Cat# 361515 |
| α-Cyano-4-hydroxycinnamic acid (4-CIN) | Sigma-Aldrich | Cat# 476870 |
| Isoflurane (inhalation anesthetic) | Baxter | Cat# ISO-250 |
| Urethane | Sigma-Aldrich | Cat# U2500-500G |
| Sucrose | JUNSEI | Cat# 31365-0350 |
| Phosphate-buffered saline (PBS, 10x), pH 7.4 | Gibco, Thermo Fisher Scientific | Cat# 70011044 |
| Triton X-100 | Sigma | Cat# T8787 |
| Methanol | Merk | Cat# 106009 |
| Normal goat serum (blocking reagent, 10 mL) | Jackson ImmunoResearch | Cat# 005-000-121 |
| Bovine serum albumin (BSA) | Sigma-Aldrich | Cat# A9647 |
| Critical commercial assays | | |
| Glycogen Colorimetric/Fluorometric Assay Kit | Abcam | Cat# ab65620 |
| SuperSignal™ West Pico PLUS substrate | Thermo Fisher | Cat# 34580 |
| TRIzol™ Reagent | Invitrogen | Cat# 15596026 |
| TOPscript™ RT DryMIX | Enzynomics | Cat# RT200 |
| Pierce BCA Protein Assay Kit | Thermo Fisher | Cat# 23225 |

| | | |
|---|---|---|
| Deposited data | | |
| RNA-seq raw data | Zhang *et al.*, 2022, Bioengineered | Supplementary Data; DOI 10.1080/21655979.2021.2021710 |
| Experimental models: Cell lines | | |
| None | | |
| Experimental models: Organisms/strains | | |
| Mouse: C57BL/6J | DooYeol Biotech | N/A |
| Mouse: B6. Cg-Tg.Gfap-cre.77.6Mvs/2J | Jackson Laboratory | N/A |
| Mouse: B6.129-Rpl22tm1.1Psam/J | Jackson Laboratory | N/A |
| Oligonucleotides | | |
| Genotyping primers (GFAP-Cre-Fw/Rv, RiboTag-Fw/Rv) | This paper | Sequences provided in Methods |
| qPCR primers (Ppp1r3c, Gys1, Phka2, Pygb, Hif1a, Ldha, Ldhb, Mct4, Hk1, Pfk1, Pkm, Pdha1) | This paper | Sequences provided in Methods |
| Recombinant DNA | | |
| None | | |
| Software and algorithms | | |
| GraphPad Prism (v9) | GraphPad Software | RRID:SCR_002798 |
| ImageJ | ImageJ | RRID:SCR_003070 |
| Thermo Xcalibur | Thermo Fisher Scientific | RRID: SCR_014593 |
| Chenomx NMR Suite | Chenomx | RRID:SCR_014682 |
| MetaboAnalyst v6.0 | Xia Lab | http://www.metaboanalyst.ca |
| GalaxyDockWeb | Seok Lab | https://galaxy.seoklab.org |
| GalaxySagittarius-AF | Seok Lab | https://galaxy.seoklab.org |
| CBDock2 | Yang Chao Lab | https://cadd.labshare.cn/cb-dock2/index.php |
| DrugRep | Yang Chao Lab | http://cao.labshare.cn/drugrep/ |
| ChimeraX | UCSF | RRID:SCR_015872 |
| R 4.3.1 | R core Team | https://www.r-project.org/ |
| ZEN | Zeiss | 3.9 |
| Biorender | Biorender | https://www.biorender.com) |
| Other | | |
| Von Frey hairs - Touch-Test Sensory Evaluators (20-piece kit) | Stoelting | Cat# 58011 |
| Injection syringe, 10 μL | Hamilton Company | Cat# 80300 |

For each ligand's table, scores were normalized within-sheet to place components on a common 0–1 scale:

$$pred_{norm} = \frac{p - \min(p)}{\max(p) - \min(p)}, \quad dock_{norm} = \frac{(-d) - \min{(-d)}}{\max(-d) - \min{(-d)}}$$

Where p is Predock and d is the docking energy (sign-flipped so that larger is better). The platform's composite score was then reconstructed by ordinary least squares (OLS) per sheet as:

$$Score(base_{recalc}) = a \cdot pred_{norm} + b \cdot dock_{norm} + c$$

With coefficients (a,b,c) fit to the sheet data ($R^2 \approx 1$ for both ligands).

Because both compounds are glycogen phosphorylase (GP) inhibitors, we applied a small heuristic class adjustment based on target class (UniProt mapping). The Final priority score used for ranking and plotting was:

$$Final\ priority\ score = Score(base_{recal}) + class_{adj}$$

With additive adjustments: GP +0.05, kinase −0.08, serine protease −0.10, metalloprotease −0.10, HSP90 −0.05, carbonic anhydrase −0.06, nuclear receptor −0.04, albumin −0.06, unknown 0. GP isoforms were defined as PYGL (P06737), PYGM (P11217), and PYGB (P11216).

Targets were rank-ordered by the Final priority score (y-axis; Figures S5D–E). GP isoforms were highlighted in red, and insets report raw Predock score and Docking energy (GalaxyDock BP2; Figures S5D-E) for the top GP hits.

### Supplementary Figures

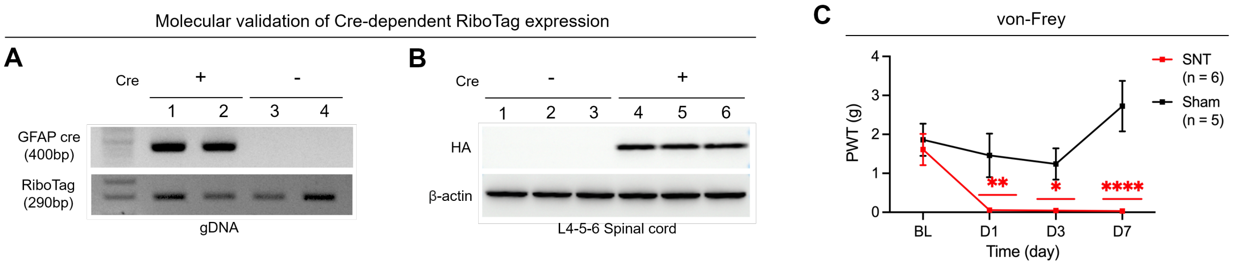


**Figure S1. GFAP Cre$^{+}$ x RPL22HA/HA RiboTag mice model validation.**

(A) PCR genotyping of tail-biopsy gDNA to detect the Cre transgene (400 bp) and the RiboTag (RPL22^HA) allele (290 bp). Lanes 1–2: Cre$^{+}$; lanes 3–4: Cre$^{-}$.

(B) Western blot of L4–L6 spinal cord lysates probed with anti-HA (top) to detect HA-tagged RPL22 and anti-β-actin (bottom) as loading control. Cre$^{+}$ mice (lanes 1–2) show robust HA signal; Cre$^{-}$ mice (lanes 3–6) do not.

(C) von-Frey test of neuropathic pain model. Sham-operated (black, n = 5) and SNT (red, n = 6) mice, measured at baseline (BL) and days 1, 3, 7 post-surgery. **P = 0.0011 (D1 Saline + SNT vs. Saline + Sham), *P = 0.0129 (D3 Saline + SNT vs. Saline + Sham), ****P < 0.0001 (D7 Saline + SNT vs. Saline + Sham)

Data are represented as the mean ± SEM; *P < 0.05, **P < 0.01, ***P < 0.001, ****P < 0.0001; Two-way ANOVA followed by Bonferroni’s multiple comparisons test (C).

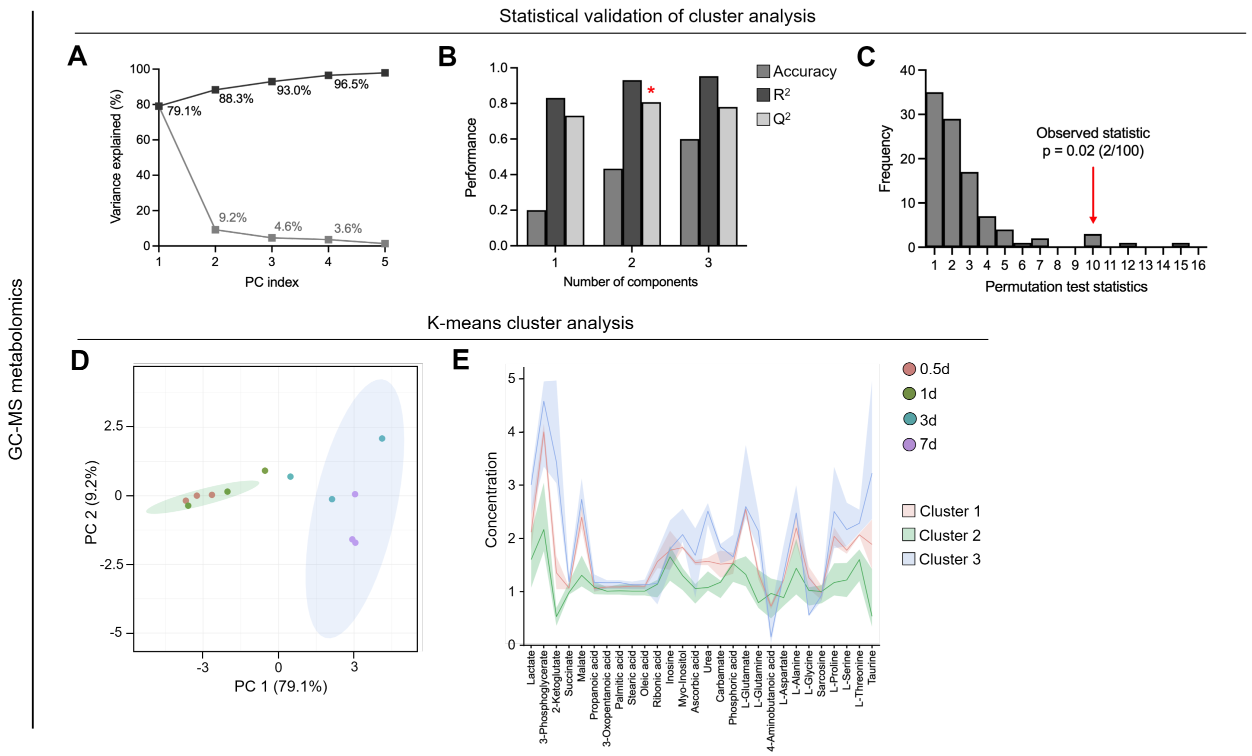


**Figure S2. Statistical analysis of GC-MS metabolomics. (Related in Fig 2.)**

(A–C) Statistical validation of cluster analysis.

(A) Cumulative variance explained by successive principal components (PCs). PC 1 accounts for 79.1 % of the total variance, with PCs 2–5 contributing 9.2 %, 4.6 %, 3.6 % and < 1 %, respectively.

(B) Cross-validation performance of the PLS-DA model as a function of the number of latent components. Bars represent classification accuracy, goodness-of-fit ($R^2$), and predictive ability ($Q^2$). A red asterisk indicates the optimal model (two components) selected for downstream analysis.

(C) Permutation test (n = 100) of the PLS-DA classification statistic. The histogram shows the distribution of test statistics obtained under random class assignments; the red arrow denotes the observed statistic (p = 0.02, 2/100 permutations ≥ observed).

(D and E) k-means cluster analysis of GC-MS metabolomics data.

(D) PCA score plot of individual samples colored by time point (0.5d: red; 1d: green; 3d: teal; 7d: purple), with 95 % confidence ellipses overlaid for the three K-means clusters identified. Axes are labeled with the percentage of variance explained by PC 1 (79.1 %) and PC 2 (9.2 %).

(E) Mean concentration profiles (± SD shading) of metabolites within each of the three clusters, plotted across time points. Cluster 1 (salmon), Cluster 2 (light green) and Cluster 3 (light blue) show distinct temporal patterns, highlighting early-, intermediate- and late-responding metabolite groups.

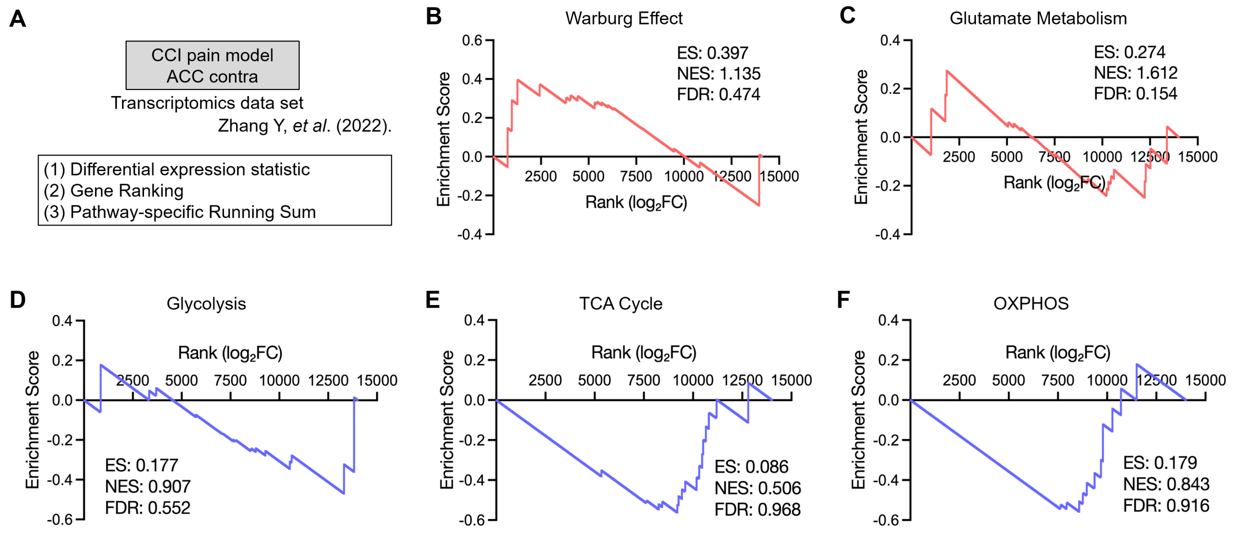


**Figure S3. ACC transcriptomics pathway analysis in chronic pain.**

(A) Experimental scheme of transcriptomics analysis in chronic constriction injury (CCI) pain model. (Time point: 7 days; fold change was normalized by Sham)

(B–F) Gene set enrichment analysis (GSEA) plot showing each pathway gene in CCI 7d (n = 3) vs. Sham (n = 3).

(B) Warburg effect metabolism GSEA plot.

(C) Glutamate metabolism GSEA plot.

(D) Glycolysis metabolism GSEA plot.

(E) TCA cycle GSEA plot.

(F) OXPHOS GSEA plot.

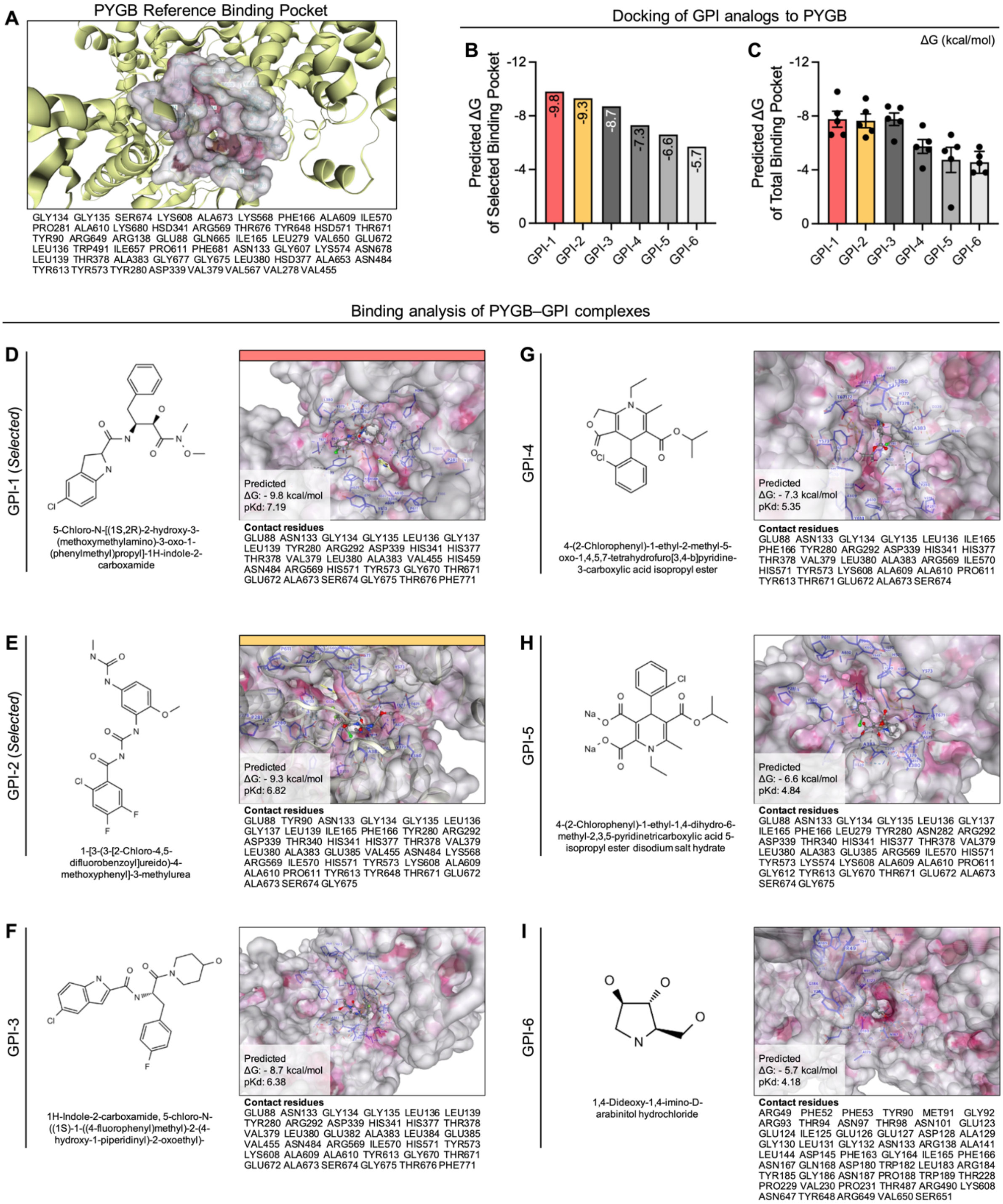


**Figure S4. Docking analysis of GPI analogs in the PYGB reference binding pocket.**

(A) Surface representation of brain glycogen phosphorylase (PYGB) showing the reference binding pocket and a docked pose of the reference GPI scaffold; pocket-lining residues are listed below.

(B–C) Predicted binding free energy (ΔG, kcal/mol) of six GPI analogs (GPI-1–GPI-6) docked to PYGB.

(B) Mean ΔG values for the selected reference pocket.

(C) Mean ΔG values for the total binding region, with individual docking poses overlaid as dots. GPI-1 and GPI-2 exhibit the lowest (most favorable) ΔG values in the selected pocket and remain among the best-scoring ligands when all poses are considered.

(D–I) Chemical structures (left) and representative docked poses (right) of GPI-1–GPI-6 within the PYGB pocket. Insets report the predicted ΔG and pKd together with the list of contact residues for each complex. GPI-1 and GPI-2 (D, E; labeled "Selected") form dense interaction networks with catalytically and structurally important residues in the pocket, including GLU88, ASN133, GLY134–137, TYR280, ASP339, HIS341, HIS377, THR378, VAL379, LEU380, ALA383, ARG569, ILE570, HIS571, TYR573, TYR613, and GLU672–SER674, and were therefore chosen as lead PYGB inhibitors for subsequent experimental evaluation.

Data are represented as the mean ± SEM.

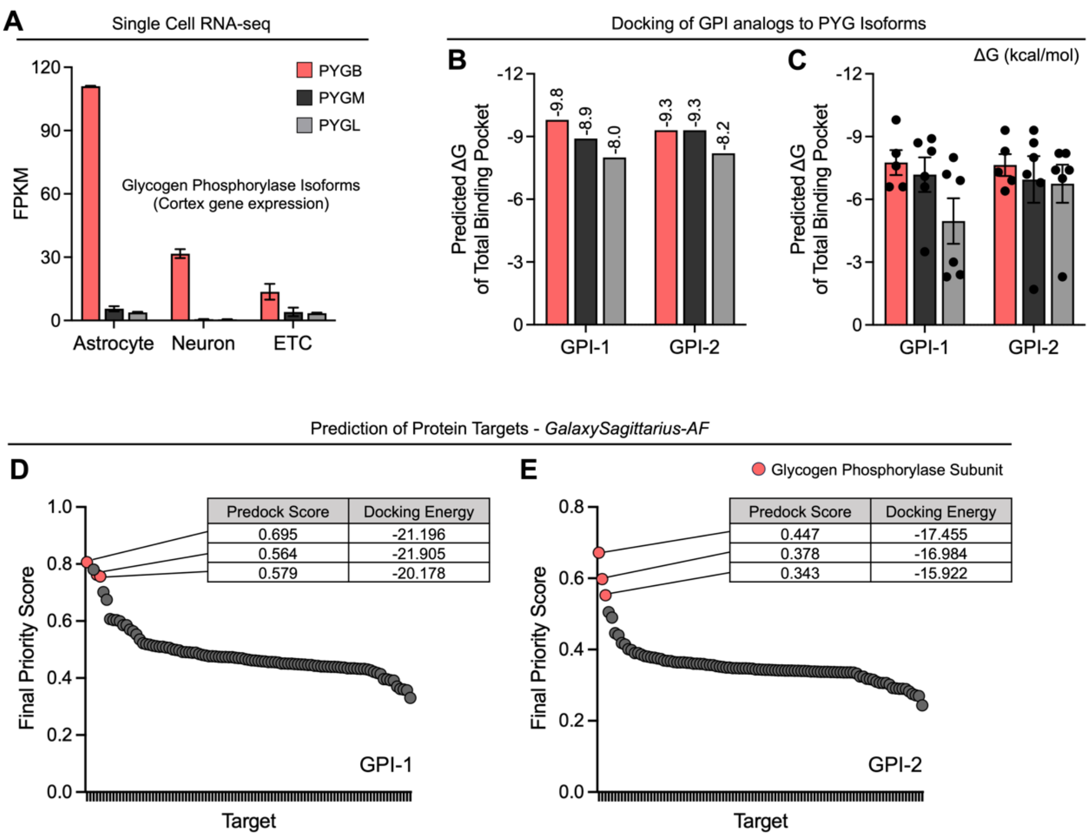


**Figure S5. Glycogen phosphorylase isoform enrichment and off-target validation.**

(A) Single-cell RNA-seq of cortex showing expression of glycogen phosphorylase isoforms. Bars indicate FPKM for PYGB (brain), PYGM (muscle), and PYGL (liver) across astrocytes, neurons, and other cell types. Data are represented as the mean ± SEM.

(B–C) Docking of GPI-1 and GPI-2 to glycogen phosphorylase isoforms. Bars show the predicted binding free energy (ΔG, kcal/mol) within the total binding pocket for PYGB, PYGM, and PYGL.

(B) Mean ΔG values for each compound–isoform pair calculated from selected docking poses.

(C) Mean ΔG values with individual docking poses overlaid as dots, illustrating that GPI-1 and GPI-2 engage the conserved catalytic pocket of all three isoforms with broadly comparable predicted affinities.

(D–E) Rank-ordered target profiling of GPI-1 and GPI-2 generated with GalaxySagittarius-AF. The y-axis shows the Final priority score and the x-axis shows each protein target (Targets n=100, ordered by rank). Red markers highlight glycogen phosphorylase isoforms among the top-ranked hits; gray points are other targets. Insets list the raw Predock score and Docking energy for the top GP hits (GalaxyDock BP2; scoring-function value). Y-axis scales are matched between panels.

(D) Rank-ordered target profiling of GPI-1.

(E) Rank-ordered target profiling of GPI-2.

Data are represented as the mean ± SEM.

**A**

MSiqDB Hallmark

| Index | Name | P-value | Adjusted p-value | Odds Ratio | Combined score |
|---|---|---|---|---|---|
| 1 | Glycolysis | 0.000001176 | 0.00001058 | 80.80 | 1103.16 |
| 2 | Hypoxia | 0.003420 | 0.01026 | 28.56 | 162.17 |

**B**

TF Perturbations Followed by Expression

| Index | Name | P-value | Adjusted p-value | Odds Ratio | Combined score |
|---|---|---|---|---|---|
| 1 | HIF1A KD Down | 0.00001589 | 0.004513 | 186.84 | 2064.50 |

**Figure S6. Enrichr analysis of qPCR expression patterns.**

(A–B) Enrichr analysis of 13 astrocyte RiboTag qPCR genes at the chronic stage (day 7).

(A) MSigDB Hallmark analysis highlighting enrichment of glycolysis and hypoxia-related pathways.

(B) "TF Perturbations Followed by Expression" library showing that HIF1A knockdown signatures are most strongly enriched, consistent with an HIF-1α–centered transcriptional program.

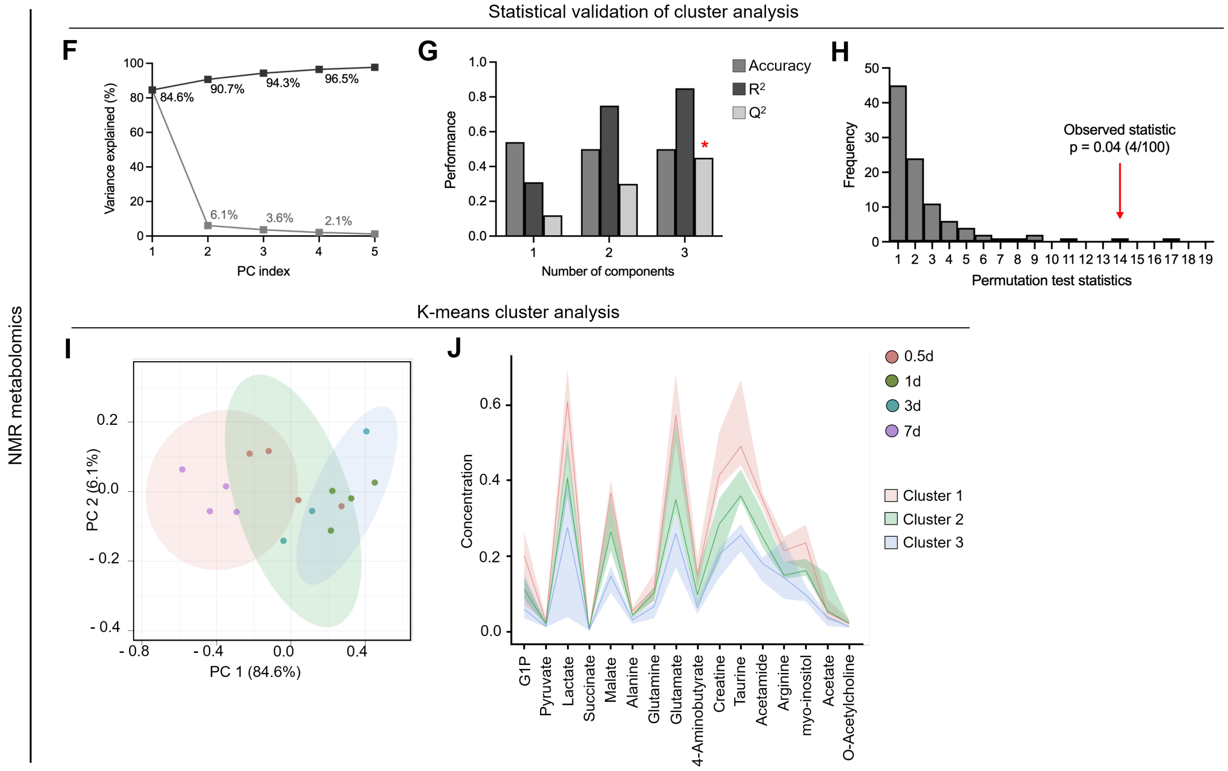


**Figure S7. Statistical analysis of NMR metabolomics. (Related in Fig 4.)**

(A–C) Statistical validation of cluster analysis.

(A) Cumulative variance explained by successive PCs. PC 1 explains 84.6% of the variance, with PCs 2–5 explaining 6.1%, 3.6%, 2.1% and < 1%, respectively.

(B) PLS-DA cross-validation metrics for one-, two- and three-component models. Light gray bars show accuracy, dark gray bars $R^2$, and white bars $Q^2$. The red asterisk marks the three-component model chosen for optimal balance of fit and prediction.

(C) Permutation testing (n = 100) of the PLS-DA statistic. The null distribution is shown as a histogram; the red arrow indicates the observed statistic (p = 0.04, 4/100 permutations ≥ observed).

(D and E) k-means cluster analysis of NMR metabolomics data.

(D) PCA score plot of all samples, colored by time point (0.5d: red; 1d: green; 3d: teal; 7d: purple) with 95% confidence ellipses for each of the three K-means clusters. PC 1 and PC 2 explain 84.6% and 6.1% of the variance, respectively.

(E) Cluster-specific mean concentration trajectories (± SD shading) of representative metabolites (listed on the abscissa). Clusters are colored as in panel D: Cluster 1 (salmon), Cluster 2 (light green) and Cluster 3 (light blue), illustrating divergent temporal responses among metabolite groups.

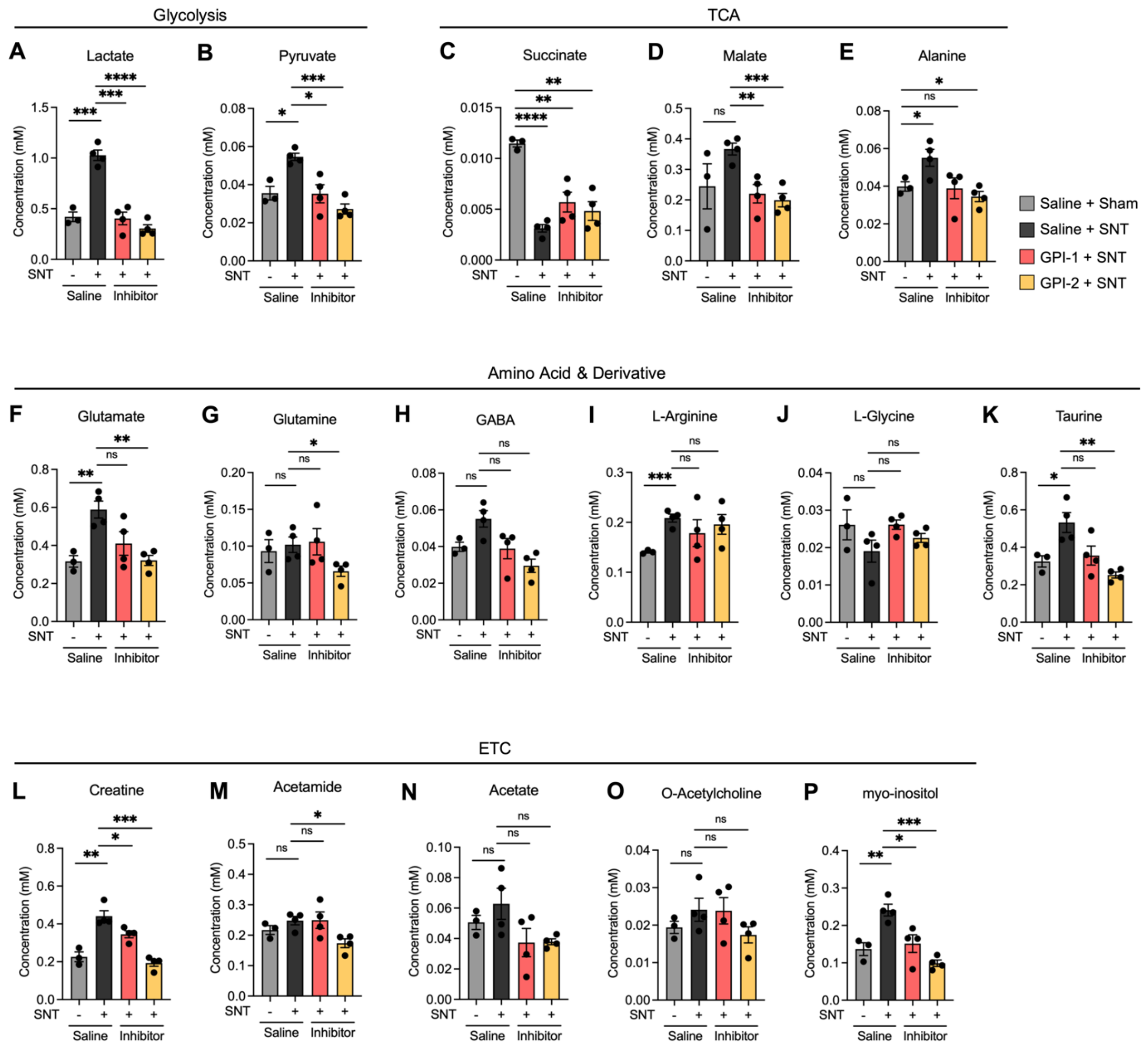


**Figure S8. Metabolite concentration data by NMR analysis. (Related in Fig 4.)**

(A) Lactate concentration of each group. ***P = 0.0003 (Saline + Sham vs Saline + SNT), ***P = 0.0003 (GPI-1 + SNT vs Saline + SNT), ****P < 0.0001 (GPI-2 + SNT vs Saline + SNT)

(B) Pyruvate concentration of each group. *P = 0.0159 (Saline + Sham vs Saline + SNT), *P = 0.0197 (GPI-1 + SNT vs Saline + SNT), ***P = 0.0003 (GPI-2 + SNT vs Saline + SNT)

(C) Succinate concentration of each group. ****P < 0.0001 (Saline + Sham vs Saline + SNT), **P = 0.0046 (GPI-1 + SNT vs Saline + SNT), **P = 0.0010 (GPI-2 + SNT vs Saline + SNT)

(D) Malate concentration of each group. **P = 0.0065 (GPI-1 + SNT vs Saline + SNT), ***P = 0.0001 (GPI-2 + SNT vs Saline + SNT)

(E) Alanine concentration of each group. *P = 0.0443 (Saline + Sham vs Saline + SNT), *P = 0.0113 (GPI-2 + SNT vs Saline + SNT)

(F) Glutamate concentration of each group. **P = 0.0056 (Saline + Sham vs Saline + SNT), **P = 0.0039 (GPI-2 + SNT vs Saline + SNT)

(G) Glutamine concentration of each group. *P = 0.0344 (GPI-2 + SNT vs Saline + SNT)

(H) GABA concentration of each group.

(I) L-Arginine concentration of each group. ***P = 0.0009 (Saline + Sham vs Saline + SNT)

(J) L-Glycine concentration of each group.

(K) Taurine concentration of each group. *P = 0.0282 (Saline + Sham vs Saline + SNT), **P = 0.0024 (GPI-2 + SNT vs Saline + SNT)

(L) Creatine concentration of each group. **P = 0.0032 (Saline + Sham vs Saline + SNT), *P = 0.0310 (GPI-1 + SNT vs Saline + SNT), ***P = 0.0003 (GPI-2 + SNT vs Saline + SNT)

(M) Acetamide concentration of each group. *P = 0.0101 (GPI-2 + SNT vs Saline + SNT)

(N) Acetate concentration of each group.

(O) O-Acetylcholine concentration of each group.

(P) Myo-inositol concentration of each group. **P = 0.0066 (Saline + Sham vs Saline + SNT), *P = 0.0197 (GPI-1 + SNT vs Saline + SNT), ***P = 0.0002 (GPI-2 + SNT vs Saline + SNT)

Data are represented as the mean ± SEM; *P < 0.05, , **P < 0.01, ***P < 0.001, ****P < 0.0001; Student's *t*-test (A-P).

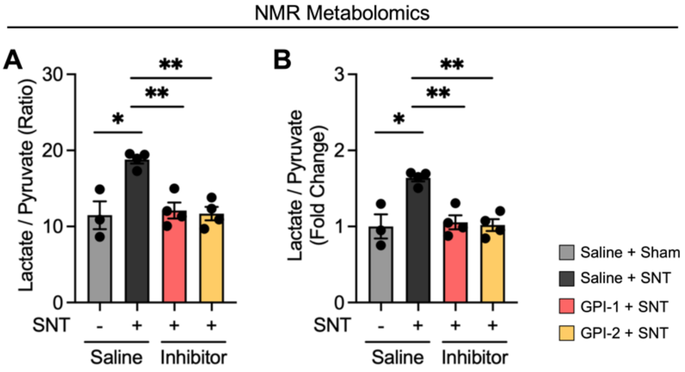


**Figure S9. Lactate/pyruvate ratio in the ACC at the chronic stage after SNT and PYGB inhibition. (Related in Fig 4.)**

(A–B) The lactate/pyruvate ratio shows a Warburg-like signature; in the Warburg effect, the lactate/pyruvate ratio is increased.

(A) Ratio of lactate/pyruvate calculated from NMR-based metabolomics data. *P = 0.0476 (Saline + Sham vs Saline + SNT), **P = 0.0034 (GPI-1 + SNT vs Saline + SNT), **P = 0.0011 (GPI-2 + SNT vs Saline + SNT).

(B) Fold change of the lactate/pyruvate ratio calculated from NMR-based metabolomics data. *P = 0.0476 (Saline + Sham vs Saline + SNT), **P = 0.0034 (GPI-1 + SNT vs Saline + SNT), **P = 0.0011 (GPI-2 + SNT vs Saline + SNT).

Data are represented as the mean ± SEM; *P < 0.05, , **P < 0.01, ***P < 0.001, ****P < 0.0001; Student's t-test (A–B).

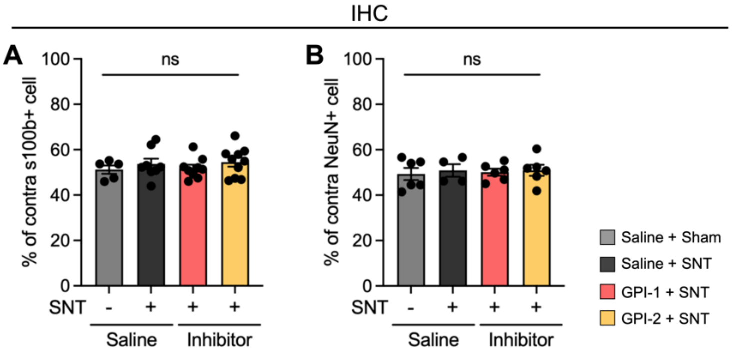


**Figure S10. Immunohistochemical assessment of astrocyte and neuronal cell density following GPI-1 and GPI-2 treatment. (Related in Fig 5.)**

(A) Ratio of s100b$^+$ cells in ACC contra area, compared with ipsi area. (Shows astrocyte cell density)

(B) Ratio of NeuN$^+$ cells in ACC contra area, compared with ipsi area. (Shows neuron cell density)

Data are represented as the mean ± SEM; *P < 0.05, , **P < 0.01, ***P < 0.001, ****P < 0.0001; Student's *t*-test (A-B).

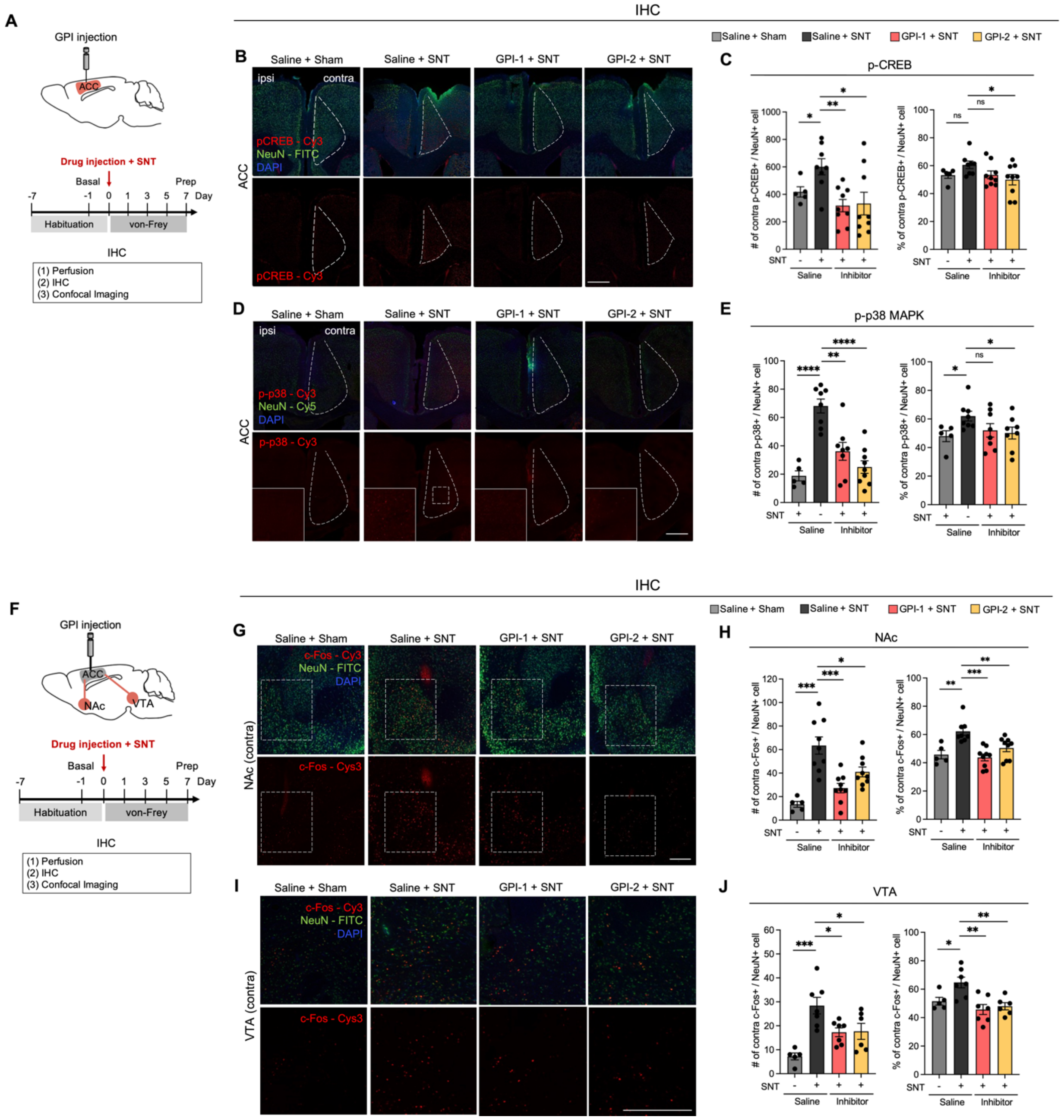


**Figure S11. Inhibition of ACC glycogenolysis decreases neuronal activation in neuropathic pain chronification. (Related in Fig 5.)**

(A) Experimental Scheme of IHC after GPI-1 and GPI-2 injection and SNT surgery.

(B and C) IHC data of p-CREB$^+$/NeuN$^+$ cells in ACC.

(B) Representative confocal images of ACC (Scale bar, 200 μm).

(C) p-CREB$^+$/NeuN$^+$ cell counting analysis data (left): Number of p-CREB$^+$/NeuN$^+$ cells in ACC contra area. *P = 0.0451 (Saline + Sham vs. Saline + SNT), **P = 0.0015 (GPI-1 + SNT vs. Saline + SNT), *P = 0.0015 (GPI-2 + SNT vs. Saline + SNT) (right): Ratio of p-CREB$^+$/NeuN$^+$ cells in ACC contra area, compared with ipsi area. *P = 0.0390 (GPI-2 + SNT vs. Saline + SNT)

(D and E) IHC data of p-p38$^+$/NeuN$^+$ cells in ACC.

(E) Representative confocal images of ACC (Scale bar, 200 μm).

(E) p-p38$^+$/NeuN$^+$ cell counting analysis data (left): Number of p-p38$^+$/NeuN$^+$ cells in ACC contra area. ****P < 0.0001 (Saline + Sham vs. Saline + SNT), **P = 0.0013 (GPI-1 + SNT vs. Saline + SNT), ****P < 0.0001 (GPI-2 + SNT vs. Saline + SNT) (right): Ratio of p-p38$^+$/NeuN$^+$ cells in ACC contra area, compared with ipsi area. *P = 0.0218 (Saline + Sham vs. Saline + SNT), *P = 0.0422 (GPI-2 + SNT vs. Saline + SNT)
(F) Experimental Scheme of IHC after GPI-1 and GPI-2 injection and SNT surgery.
(G-J) IHC data of c-Fos$^+$/NeuN$^+$ cells in NAc.
(G) Representative confocal images of NAc (Scale bar, 200 µm).
(H) IHC data of c-Fos$^+$/NeuN$^+$ cells in NAc (left): Number of c-Fos$^+$/NeuN$^+$ cells in NAc contra area. ***P = 0.0004 (Saline + Sham vs. Saline + SNT), ***P = 0.0005 (GPI-1 + SNT vs. Saline + SNT), *P = 0.0175 (GPI-2 + SNT vs. Saline + SNT) (right): Ratio of c-Fos$^+$/NeuN$^+$ cells in NAc contra area, compared with ipsi area. **P = 0.0004 (Saline + Sham vs. Saline + SNT), ***P = 0.0005 (GPI-1 + SNT vs. Saline + SNT), ***P = 0.0175 (GPI-2 + SNT vs. Saline + SNT)
(I-J) IHC data of c-Fos$^+$/NeuN$^+$ cells in VTA.
(I) Representative confocal images of VTA (Scale bar, 500 µm).
(J) IHC data of c-Fos$^+$/NeuN$^+$ cells in NAc (left): Number of c-Fos$^+$/NeuN$^+$ cells in VTA contra area. ***P = 0.0006 (Saline + Sham vs. Saline + SNT), *P = 0.0144 (GPI-1 + SNT vs. Saline + SNT), *P = 0.0484 (GPI-2 + SNT vs. Saline + SNT) (right): Ratio of c-Fos$^+$/NeuN$^+$ cells in VTA contra area, compared with ipsi area. *P = 0.0256 (Saline + Sham vs. Saline + SNT), **P = 0.0031 (GPI-1 + SNT vs. Saline + SNT), **P = 0.0045 (GPI-2 + SNT vs. Saline + SNT)
Data are represented as the mean ± SEM; *P < 0.05, **P < 0.01, ***P < 0.001, ****P < 0.0001; Student's *t*-test (C, E, H and J).

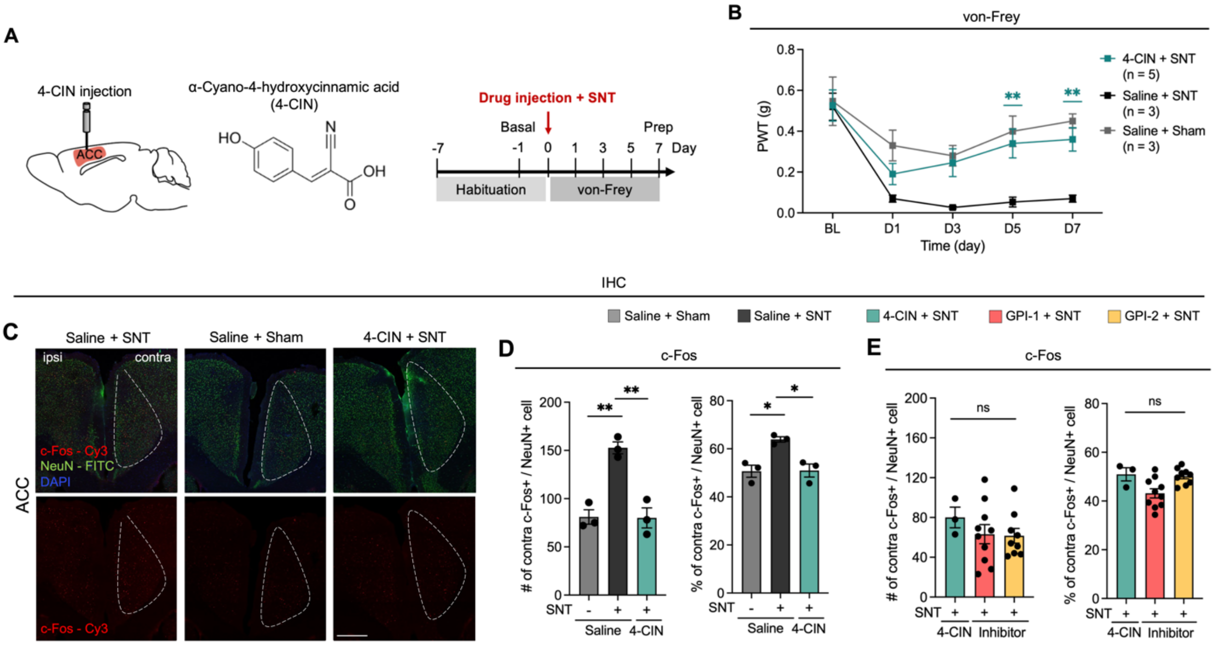


**Figure S12. ACC lactate transport mediates neuropathic pain chronification.**

(A) Experimental Scheme of IHC after 4-CIN injection and SNT surgery.

(B) von-Frey test of neuropathic pain model. **P = 0.0095 (D5 4-CIN + SNT vs. Saline + SNT), **P = 0.0087 (D7 4-CIN + SNT vs. Saline + SNT)

(C) Representative confocal images of ACC (Scale bar, 200 μm).

(D and E) IHC data of c-Fos$^{+}$/NeuN$^{+}$ cells in ACC.

(D) Saline or 4-CIN injection group analysis (left): Number of c-Fos$^{+}$/NeuN$^{+}$ cells in ACC contra area. **P = 0.0021 (Saline + Sham vs. Saline + SNT), **P = 0.0075 (4-CIN + SNT vs. Saline + SNT) (right): Ratio of c-Fos$^{+}$/NeuN$^{+}$ cells in ACC contra area, compared with ipsi area. *P = 0.0214 ((Saline + Sham vs. Saline + SNT), *P = 0.0277 (4-CIN + SNT vs. Saline + SNT).

(E) Compare IHC data with 4-CIN injection experiment (left): Number of c-Fos$^{+}$/NeuN$^{+}$ cells in ACC contra area (right): Ratio of c-Fos$^{+}$/NeuN$^{+}$ cells in ACC contra area, compared with ipsi area.

Data are represented as the mean ± SEM; *P < 0.05, **P < 0.01, ***P < 0.001, ****P < 0.0001; Two-way ANOVA-multiple comparisons (B) and Student's *t*-test (D and E).

### Methods Validations

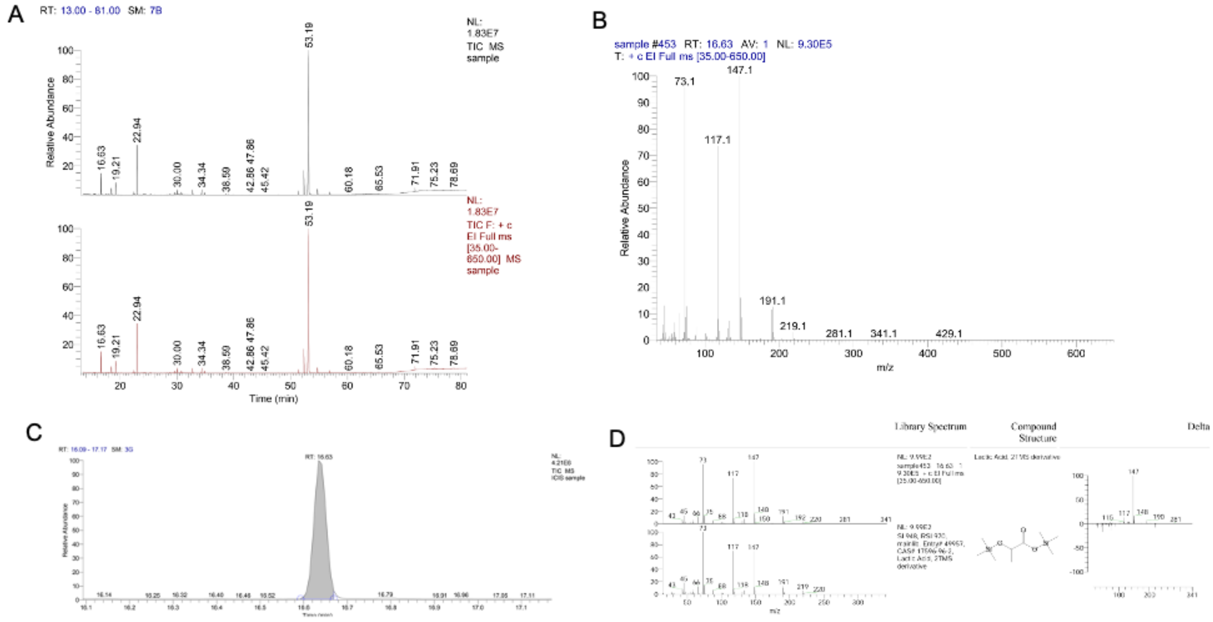


**Representative GC–MS raw data for lactate (2TMS derivative)**

(A) Representative TIC focused on the target retention-time window (RT 16.09–17.17 min); the lactate derivative elutes at RT 16.63 min (shaded/marked).

(B) Representative EI full-scan mass spectrum (m/z 35–650) at the peak apex; characteristic fragments m/z 73, 117, 147, 191, 219 are indicated.

(C) Representative mirror plot versus the NIST main library (compound: Lactic acid, 2TMS derivative); SI/RSI values as shown in the panel, with the library structure inset.

(D) Representative full-run TIC (13–81 min) providing chromatographic context with the lactate peak position marked.

Data shown are representative of the dataset used for metabolite calling.

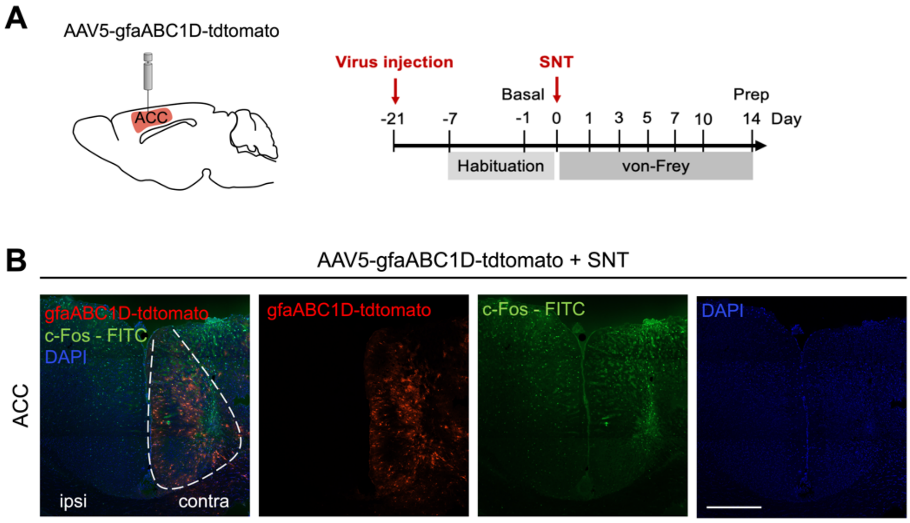


**Validation of contralateral ACC-targeted drug injection and viral expression.**

(A) Schematic of ACC-targeted infusion and schedule. AAV5-gfaABC1D-tdTomato was injected 21 d before spared nerve transection (SNT; day 0); animals underwent habituation (days −7 to −1), baseline assessment (day −1), and von Frey testing on days 1, 3, 5, 7, and 10; tissue was prepared on day 14.

(B) Representative IHC confocal data of the ACC (coronal sections) from an AAV5-gfaABC1D-tdTomato–injected, SNT-operated mouse showing tdTomato fluorescence (astrocyte-restricted gfaABC1D promoter, red), c-Fos immunoreactivity (FITC, green), and DAPI (blue). Composite (left) and single-channel images (middle/right) are shown; ipsilateral and contralateral sides are indicated. This figure validates ACC targeting and astrocyte-selective viral expression used for the pharmacological experiments. (Scale bar, 200 μm).
